# Fat1 deletion enhances Fibro-Adipogenic Differentiation and Adipogenic expansion following injury in skeletal muscle

**DOI:** 10.64898/2026.04.21.719880

**Authors:** Pierre-Antoine Ferracci, Françoise Helmbacher

## Abstract

Skeletal muscles regenerate following injury, owing not only to myogenic stem cells, but also to non-myogenic cells such as fibro-adipogenic progenitors (FAPs). Quiescent in healthy muscles, FAPs transiently proliferate in response to tissue-damage, to support myogenic repair. Aside from their pro-myogenic role in healthy muscles, FAPs are also the origin of intramuscular fibro-adipose tissue that infiltrate muscles with chronic inflammation and degeneration in various muscle pathologies. Here, we investigate how the Fat1 Cadherin, previously identified as a regulator of embryonic muscle morphogenesis, influences FAP biology during damage-induced muscle regeneration. *Fat1* expression is transiently induced in FAPs and myogenic cells after muscle damage. We found that mesenchyme-specific *Fat1* ablation leads to increased fibro-adipogenic infiltrations following glycerol injury, while minimally affecting myogenic repair. Using an inducible Pdgfra-cre/ERT line, we further demonstrated that *Fat1* restricts FAP adipogenic differentiation through both cell-autonomous and non-cell-autonomous mechanisms. These findings identify *Fat1* as a novel regulator of FAP biology, essential for limiting FAP differentiation and the development of fibro-fatty infiltrations after muscle injury.

## Introduction

Skeletal muscles have a remarkable capacity to regenerate after acute tissue lesions. This regenerative capacity is permitted not only by myogenic stem cells (satellite cells), which can replace damaged fibers, but also by an array of non-myogenic muscle-resident cells constituting a supportive niche (Biferali *et al*, 2019; Helmbacher & Stricker, 2020; Relaix & Zammit, 2012; Theret *et al*, 2021). Among those, fibro-adipogenic progenitors (FAPs) are muscle-resident mesenchymal cells that have gained considerable interest since their initial description (Joe *et al*, 2010; Uezumi *et al*, 2010). While they do not directly contribute to the myogenic lineage, they are necessary for skeletal muscle mass maintenance, homeostasis and regeneration (Wosczyna *et al*, 2019). Mostly quiescent in healthy muscles, FAPs proliferate in response to tissue lesions to support myogenic repair and ECM remodelling (Joe *et al*., 2010; Uezumi *et al*., 2010), thus massively invading interstitial space in the lesioned area, and leading to a state of transient fibrosis (Joe *et al*., 2010). In this phase, they secrete various factors, ECM components and ECM-associated proteins that promote myogenic repair (Helmbacher & Stricker, 2020; Joe *et al*., 2010; Lukjanenko *et al*, 2019; Theret *et al*., 2021; Uezumi *et al*., 2010; Uezumi *et al*, 2021). Blocking FAP expansion by pharmacological inhibition (Fiore *et al*, 2016), by genetic ablation experiments (Wosczyna *et al*., 2019), or by altering their secretome (Uezumi *et al*., 2021), compromises the efficiency of muscle regeneration, and leads to muscle atrophy and depletion of the satellite cells pool. FAPs also orchestrate the dynamic contributions of inflammatory cells to muscle repair, by promoting the clearance of debris of degenerating myofibers, and subsequently by secreting cytokines promoting the pro- to anti-inflammatory switch in macrophage phenotype (Heredia *et al*, 2013; Lemos *et al*, 2015). Once the repair process induced by acute damage has started, FAPs are subsequently eliminated by apoptosis, triggered by signals produced by inflammatory macrophages (Lemos *et al*., 2015), thus making space for growth, fusion, and maturation of de-novo fibers.

Besides their pro-myogenic activity, FAPs are multipotent progenitors known for their inherent capacity to give rise to adipocytes, tissue-fibroblasts/myofibroblasts, osteoblasts, and a few other stromal derivatives (Eisner *et al*, 2020; Santini *et al*, 2020; Uezumi *et al*, 2011). While naturally-occurring differentiation is minimal in steady-state healthy muscles, ectopic differentiation can occur in regenerating muscle, but is strictly limited and transient in healthy context (Lukjanenko *et al*, 2013). Instead, ectopic differentiation is unleashed in the context of a number of muscle pathologies, including Duchenne Muscular Dystrophy (DMD) (Contreras *et al*, 2016; Mozzetta *et al*, 2013; Trensz *et al*, 2010; Uezumi *et al*., 2011), Amyotrophic lateral sclerosis (ALS) (Gonzalez *et al*, 2017; Madaro *et al*, 2018), Facio-scapulo-humeral dystrophy (FSHD) (Bosnakovski *et al*, 2017; Dandapat *et al*, 2014), or limb girdle muscular dystrophy (LGMD) (Hogarth *et al*, 2019). Progression of this pathological differentiation leads to the replacement of the muscle mass by fibrosis and adipose infiltrations, commonly referred to as fibro-fatty infiltrations, or as inter- and intra-muscular adipose tissue (IMAT), jointly referred to as myosteatosis (Correa-de-Araujo *et al*, 2020; Flores-Opazo *et al*, 2024). The extent of myosteatosis in diseased muscles, visualized in patients by magnetic resonance imaging (MRI) (Correa-de-Araujo *et al*., 2020; Godi *et al*, 2016; Kan *et al*, 2009; Lareau-Trudel *et al*, 2015), is inversely correlated with regenerative capacities (Norris *et al*, 2024; Norris *et al*, 2025), implying that these fibro-fatty infiltrates compromise the efficacy of myogenic regeneration and fiber growth, directly or through their effects on the physical properties of muscle. Thus, aside from their respective genetic causes, a common feature of chronic muscle pathologies is that they interfere with a gatekeeping system that allows maintaining FAP quiescence, suppressing excessive FAP differentiation, ultimately eliminating excess FAPs and their differentiation byproducts in healthy muscles. Despite being under intense scrutiny by the biomedical community since the discovery that FAPs are the cell type of origin of fibro-fatty deposition, the mechanisms involved in suppressing FAP differentiation are still poorly understood. Thus, identifying novel regulatory mechanisms limiting the formation of fibro-fatty tissue represents a key unmet medical need to delay disease progression.

In the present study, we investigated the role exerted by the Fat1 Cadherin in FAPs during skeletal muscle regeneration induced by acute lesions. Fat-like Cadherins are adhesion molecules that regulate coordinated cell behaviors such as planar cell polarity in epithelia (Bosveld *et al*, 2012; Sharma & McNeill, 2013a, b), polarized cell movements (Caruso *et al*, 2013; Zakaria *et al*, 2014), and oriented cell divisions (Mao *et al*, 2016; Saburi *et al*, 2008), and that control tissue growth via the Hippo pathway (Cho *et al*, 2006; Lawrence & Casal, 2013; Matis & Axelrod, 2013). Among them, we previously identified *Fat1* as a regulator of neuromuscular morphogenesis during mouse development (Caruso *et al*., 2013; Helmbacher, 2018). *Fat1* influences the growth and the shape of subsets of muscles in the embryo, through complementary activities in different cell types including myogenic cells, motor neurons and muscle-associated mesenchymal cells (Helmbacher, 2018). In humans, alterations of *FAT1* expression were observed in FSHD patients (Caruso *et al*., 2013; Mariot *et al*, 2015). Furthermore, pathological *FAT1* variants disrupting its splicing or predicted to alter its functions were identified in patients with FSHD-like symptoms in absence (Puppo *et al*, 2015), but also in association with (Park *et al*, 2018) traditional FSHD-causing mutations, implicating *FAT1* as a putative modifier of FSHD severity (Caruso *et al*., 2013; Caruso *et al*, 2022; Mariot *et al*., 2015; Puppo *et al*., 2015). Guided by the observation that mesenchymal *Fat1* activity represented the predominant component of its role in muscle morphogenesis during development, and by the fact that adult FAPs derive from the developing connective tissue (Stumm *et al*, 2018; Vallecillo-Garcia *et al*, 2017), we enquired whether *Fat1* signaling might be required in adult FAPs to ensure proper muscle regeneration and homeostasis.

We found that *Fat1* deletion in the mesenchymal lineage enhanced the fibro-adipogenic expansion induced by glycerol injury in skeletal muscle, with minimal impact on myogenic repair. Owing to an inducible CRE line leading to low amounts of recombined FAPs, we found *Fat1* is required to prevent FAP differentiation, and that loss of *Fat1* in FAPs enhances intramuscular adipogenesis and IMAT deposition both cell-autonomously and non-cell-autonomously. Distinguishing phenotypes by sex uncovered that enhancement of adipose infiltrations in mutants was more robust in females, whereas enhancement of transient fibrosis was only observed in males. Futhermore, the cell-autonomous and non-cell autonomous components of the adipogenesis phenotype were unequally associated with sex. Altogether, these data identify *Fat1* as a modulator of FAP differentiation, fate choice, and homeostasis, required to limit the expansion of glycerol-injury-induced adipose infiltrates, with sex-associated features that might guide future assessment of patient data.

## Results

### Fat1 expression is induced by skeletal muscle injury in subsets of FAPs and myogenic cells

*Fat1* is broadly expressed in the musculo-skeletal system during development, encompassing myogenic progenitors and their derivatives, muscle-associated connective tissues, as well as tendons, bones and cartilage (Caruso *et al*., 2013; Helmbacher, 2018; Mariot *et al*., 2015; Smith *et al*, 2007). *Fat1* expression levels vary not only between tissue types, but also between muscle subtypes, with distinct intensities of *Fat1^LacZ^* signal detected in different muscles at E13.5 (Caruso *et al*., 2013; Mariot *et al*., 2015). While *Fat1* expression is still relatively high at early postnatal stages, it undergoes a major drop in expression level at adult stages (Mariot *et al*., 2015). Thus, *Fat1* expression levels are highest at stages of active developmental and postnatal myogenesis, whereas our studies in humans and mice illustrated that in homeostatic adult muscles, *Fat1* dosage, although lowered, keeps distinguishing several muscle groups (Mariot *et al*., 2015).

Given the strong impact that deletion of *Fat1* activities in myogenic progenitors and mesenchymal cells exerts on muscle development (Caruso *et al*., 2013; Helmbacher, 2018), and given that *Fat1* expression levels in adult muscles are considerably lower than in embryonic, fetal and juvenile muscle (Mariot *et al*., 2015), we asked whether *Fat1* expression might be reactivated during injury-induced muscle regeneration, and whether it might play functions analogous to those played during development. The *Fat1^LacZ^* mutant allele represents a convenient tool to further monitor *Fat1* expression, and to study its distribution in various tissue-types (Caruso *et al*., 2013; Helmbacher, 2018, 2022). We exposed young adult mice (2-6 months) carrying the *Fat1^LacZ^* allele to muscle injury, triggered by Cardiotoxin (Figure 1B, C, Figure S2) or by glycerol injections (Figure 1C, D-F, Figure S1). We injured either the *tibialis anterior* (TA, Figure 1B), a classical site for the study of muscle regeneration, or the *triceps brachii* (TB, Figure 1D-F), a humeral muscle belonging to the panoply of muscles affected in FSHD patients at early disease stages, which also expresses high levels of *Fat1* during muscle development (Helmbacher, 2018; Mariot *et al*., 2015). *Fat1^LacZ^* expression was monitored at 5 and 7 days post-injury (dpi), by visualizing β-galactosidase activity with the substrate salmon gal (Figure 1B,D, Figure S2B), or by immunohistochemistry with anti-β-galactosidase antibodies combined with markers of distinct cell types (Figure 1, Figures S1, S2).

**Figure 1:**
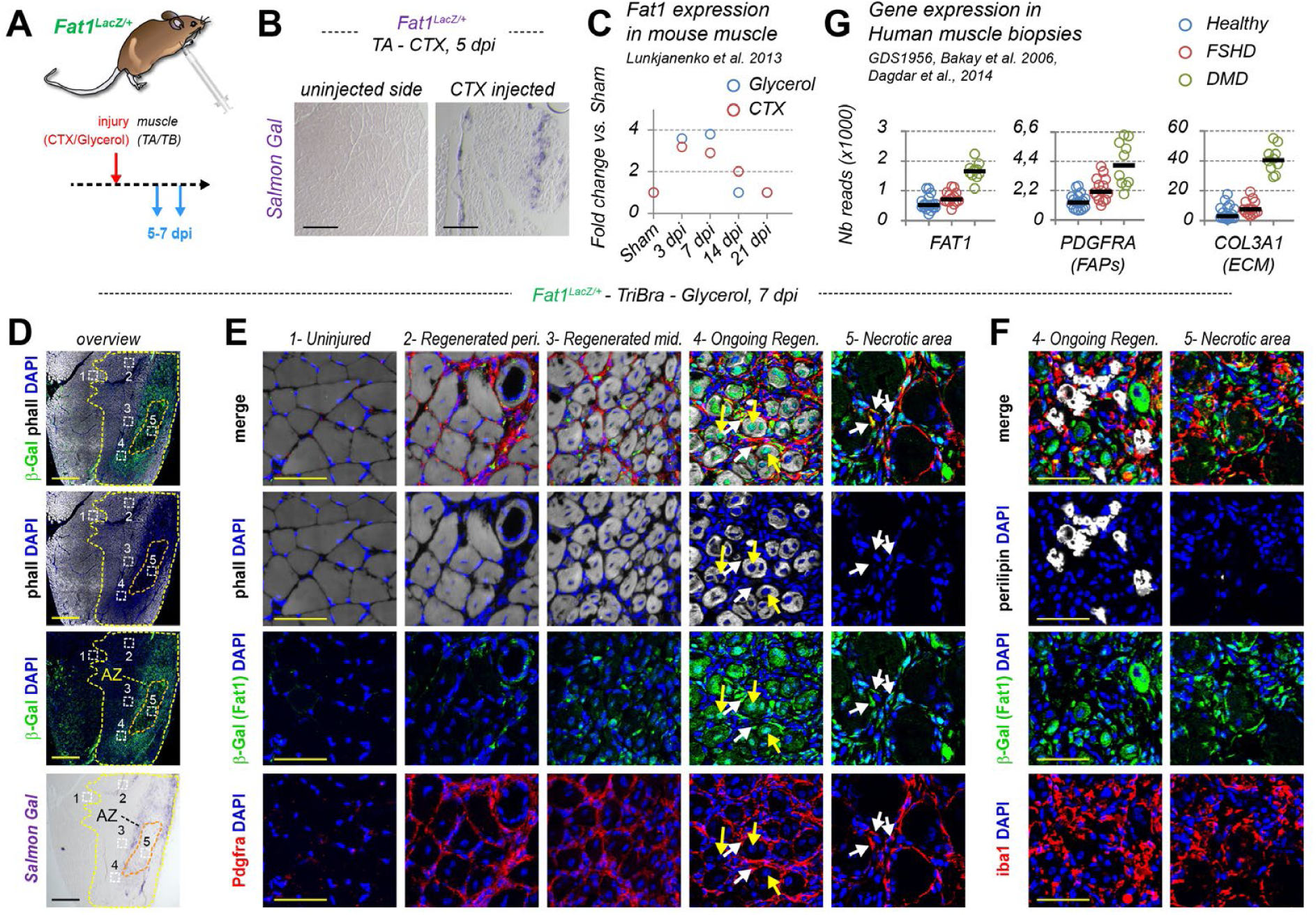
Fat1 expression is transiently induced in FAPs and myogenic cells in regenerating muscle after injury. (A) Scheme of the experiment, in which adult *Fat1^LacZ/+^* mice were injured either with glycerol injected in the Triceps Brachii muscle, or with Cardiotoxin (CTX) injected in the Tibialis anterior muscle, and the injured muscles were collected respectively at 7 or 5 days post-injury for histological analyses. (B) Salmon Gal staining of muscle cross-sections from the lesion side or the uninjected side, showing that *Fat1^LacZ/+^*expression is undetectable in the uninjured adult muscle, but induced by the CTX injury. (C) Plots of the evolution of Fat1 mRNA levels in mouse muscle after Glycerol or CTX injury at the indicated stages. Transcriptomic data were extracted from Lukjanenko et al., 2013. (D, E, F) Expression of *Fat1^LacZ^* after glycerol injury at 7 dpi was visualized by salmon Gal staining (purple, D) or by immunohistochemistry with anti-beta-galactosidase antibodies (green), combined with phalloiding (white), and antibodies against Pdgfra (red, E), or against iba1 (red, F). Images in (D) represent vues of the entire muscle cross-section, while panels in (E, F) are high magnification images of areas highlighted by dotted squares in (D). The position of high magnification panels is indicated in (D) by numbers (1- uninjured; 2- Regenerated peripheral; 3- regenerated medial; 4- ongoing regeneration; 5- necrotic area), and are placed relative to the outer limit of the lesion area (transition between fibers with peripheral and central nuclei), and to the limits of the necrotic area (where degenerating fibers exhibit reduced or absent phalloidin staining). The active zone (AZ) is the area at the transition between degenerating and regenerated fibers, where satellite cells and FAPs are mobilized to start the myogenic repair processed. While Pdgfra labels FAPs throughout the lesion, *Fat1^LacZ^* expression levels are highest in the active zone with ongoing regeneration, and in cells surrounding degenerating fibers in the necrotic area, including FAPs and a few macrophages. (G) Comparison of levels of FAT1, PDGFRA and COL3A1 levels in human muscle biopsies from healthy patients or patients with DMD or FSHD, extracted from GDS1956 Gene expression dataset from Bakay et al. 2006, Dagdar et al., 2014). Scalebars: (B) 250 µm; (D) 500 µm; (E, F) 50 µm.

*Fat1^LacZ^* expression was strongly induced in the lesion area after both cardiotoxin (5dpi) and glycerol (7dpi) injuries (Figure 1B,D, Figure S1), whereas β-galactosidase activity or protein levels were below detection levels in unaffected ipsilateral muscle areas or contralateral uninjured muscles. These observations are consistent with previously published transcriptome data from regenerating mouse muscle (Figure 1C, data from (Lukjanenko *et al*., 2013)). *Fat1^LacZ^* expression was highest at the interface between the central degenerating area (containing degenerating/necrotic fibers, or ghosts of degenerated fibers) and the more peripheral area occupied by new fibers (recognizable by centrally located nuclei). This interface corresponds to the region of active regenerative myogenesis (Figure 1D, E), where β-galactosidase was detected in the youngest de-novo produced myofibers (yellow arrows in Figure 1E), and in interstitial mesenchymal cells expressing PDGFRA, co-expression being highest in the areas with ongoing regeneration and surrounding necrotic fibers (white arrows in Figure 1E). Instead, *Fat1^LacZ^* expression was relatively lower in FAPs surrounding regenerated myofibers in the periphery of the lesioned area where regeneration is more advanced and de-novo fiber diameter is larger, indicating that *Fat1* induction is transient (compare crops 2-3 and 4-5 in Figure 1E). The resulting gradient of *Fat1* expression levels was most obvious when following staining intensities along a band spanning from the uninjured area to the center of the necrotic area (Figure S1B, C, D). This contrasts with levels of Pdgfra, which are elevated throughout the lesion area compared to the uninjured area, with a progressive increase in intensity between peripheral and central FAPs (Figure 1E, Figure S1C, D), such that Pdgfra levels are higher than those of Fat1 in peripheral FAPs, and lower in central FAPs (Figure S1D). *Fat1^LacZ^* expression was also low in PAX7-expressing satellite cells (Figure S2C), and present in small subsets of macrophages (Figure 1F and Figure S2E). This topological gradient also matches the evolution over time of *Fat1* RNA levels at later timepoints after muscle injury in published datasets (Figure 1C, (Lukjanenko *et al*., 2013)), with a sharp increase at 5 and 7 dpi, followed by a progressive return to baseline levels at 14 and 21 dpi. In GEO transcriptome datasets GDS1956 from references (Bakay *et al*, 2006; Dadgar *et al*, 2014), *FAT1* levels are elevated in both DMD and FSHD muscle biopsies vs. controls (Figure 1G), correlating with RNA levels of *PDGFRA* and *COL3A1*, hence with the degree of FAP amplification and fibrosis in these samples. Overall, these data support the possibility that Fat1 signaling might play a role during skeletal muscle regeneration, acting predominantly in the myogenic and mesenchymal lineages.

Since *Fat1* was not uniformly detected across all FAPs in the lesion area, we wondered if its expression was particularly enriched in a specific FAPs subtype or associated with a specific regenerative-like state. Multiple recent single cell RNAseq (scRNA-seq) studies carried out on mouse muscle, including both homeostatic and post-injury conditions, have illustrated the remarkable molecular heterogeneity among FAP subtypes (Collins & Kardon, 2021; De Micheli *et al*, 2020; Oprescu *et al*, 2020). These datasets have been conveniently integrated in the joint scMuscle database (McKellar *et al*, 2021), providing a valuable resource for comparative analyses. We extracted from this scMuscle dataset the scRNAseq expression levels of *Fat1* alongside those of established markers of distinct FAP subsets (Figure 2). These included *Pdgfra*, a common marker for all stromal cells, and selected subset-specific markers, such as *Hic1*, *Pi16*, *Mme*, or *TnC*. Consistent with our observations, *Fat1* RNA was detected in the FAP cluster, in Smooth muscle cells, in scattered cells among the myoblast and myogenic stem cell clusters (Figure 2A,B). Focusing on FAPs, we color-converted *Fat1* in green and the other markers in red to examine their overlap (Figure 2D). This highlighted that although *Fat1* levels are not uniform among *Pdgfra*-expressing FAPs, *Fat1* expression was not restricted to a specific FAP subtype, but displayed instead a salt-and-pepper distribution throughout all subtypes. Consistent with the transient increase of *Fat1* expression after injury, high *Fat1* levels were detected in a subgroup labelled as “pro-remodelling” FAPs, corresponding to a FAP subtype most abundant at 5 dpi (De Micheli *et al*., 2020; McKellar *et al*., 2021), and characterized by high Tenascin C (TnC) expression (Figure 2D). TnC is known as a tenocyte marker in developing and adult tendons (Edom-Vovard *et al*, 2002; Kardon, 1998), but its injury-induced expression in pro-remodelling FAP is characteristics of its activity as damaged-induced molecular pattern (Midwood *et al*, 2009; Piccinini & Midwood, 2010).

**Figure 2:**
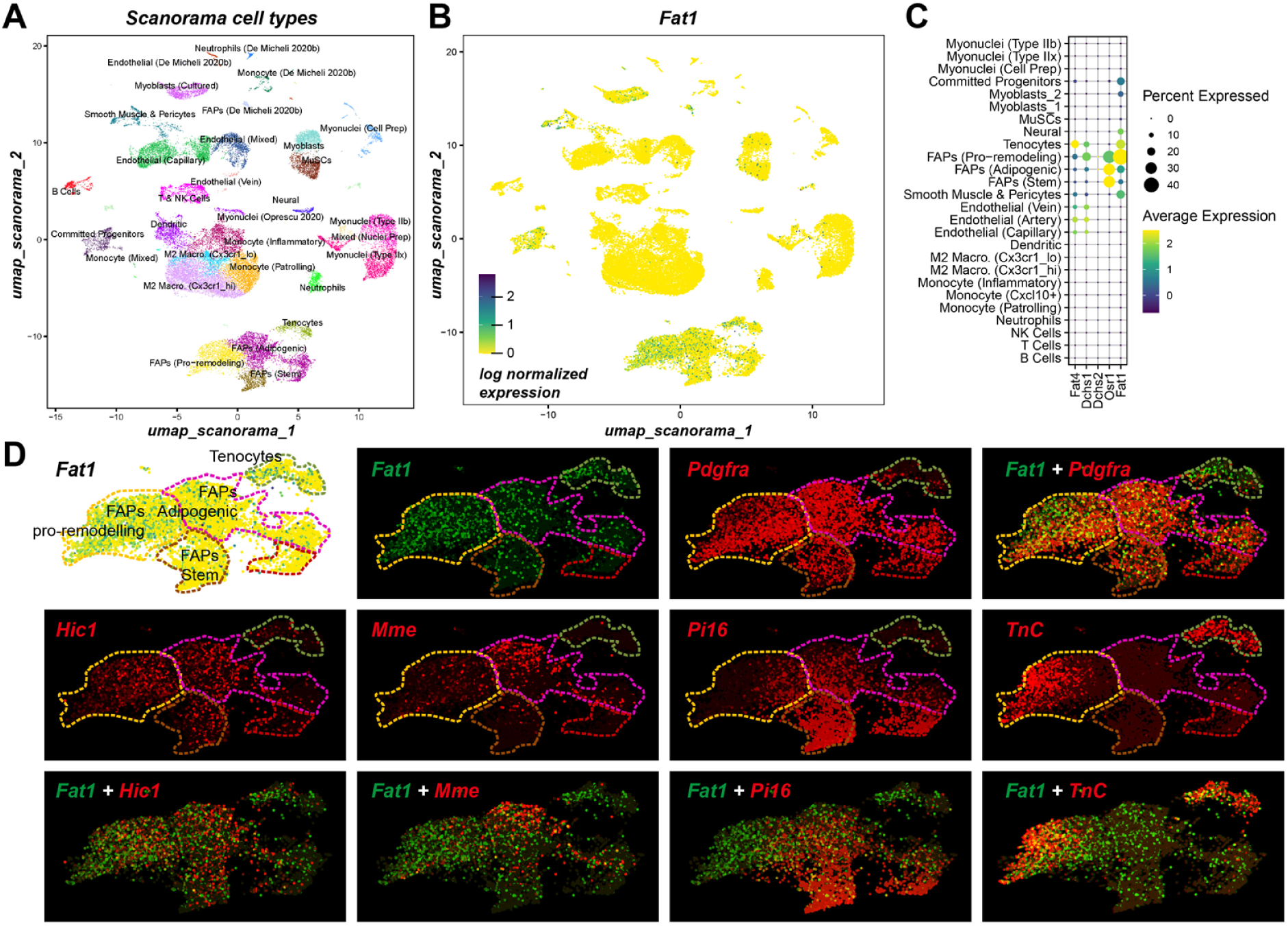
Fat1 expression in published mouse muscle ScRNAseq datasets: Distinguishing FAP subsets. (A, **B)** Image showing the umap scanorama plot with cell type annotations (A), or with Fat1 expression (B) from the ScMuscle web interface from McKellar et al (McKellar *et al*., 2021), which integrated ScRNAseq data from several published sources focusing on healthy or injured mouse muscle. In (B) the cell cluster with highest Fat1 levels include FAPs, also shown magnified in (D). **(C)** Dot plot (generated in scMuscle) showing percent of cells from each annotated cell type, and mRNA levels (colors) for Fat1, Fat4, Dchs1, Dchs2 and Osr1. **(D)** Umap Scanorama plots zooming on FAPs for Fat1, and through a color conversion, its overlap with several other fibro-adipogenic progenitors or Mesenchymal stromal cell markers (converted to red, whereas *Fat1* is converted to green), including *Pdgfra*, *Hic1*, *Mme*, *Pi16*, and *TnC*. While *Fat1*-expressing cells are detected among all FAP subtypes, the highest percentage of expressing cells and levels match with the subset expressing *TnC*, corresponding (according to (McKellar *et al*., 2021)) the FAP pro-remodelling subtype and to tenocytes.

### Impact of mesenchymal Fat1 deletion on glycerol-induced muscle regeneration and fibro-adipogenic infiltration

Guided by *Fat1* expression in pro-remodelling FAPs, and by our work dissecting how tissue-specific *Fat1* activities control muscle development, we focused on the mesenchymal lineage (Helmbacher, 2018). To explore the role played by *Fat1* in muscle-resident mesenchymal cells, we induced muscle injury in a cohort of adult *Fat1^Prx1^* and control mice. We explored this in the context of glycerol-induced lesions, a model permissive to regeneration, while also leading to some persistent fibrosis and adipose infiltrations (Lukjanenko *et al*., 2013; Norris *et al*., 2024; Pisani *et al*, 2010). We chose to perform these experiments in the triceps brachii (medial), a muscle with high *FAT1* levels (Caruso *et al*., 2013; Mariot *et al*., 2015), that belongs to the clinical map of early disease stages in FSHD patients (Mariot *et al*., 2015), rather than in the tibialis anterior muscle typically used in most regeneration studies. The Triceps Brachii (TB) muscle is also large enough for the injected myotoxic compound (i.e., glycerol) not to damage the whole muscle mass, preserving fibers in the periphery of the lesion (Figure 1E,F, and Figure 3D).

**Figure 3:**
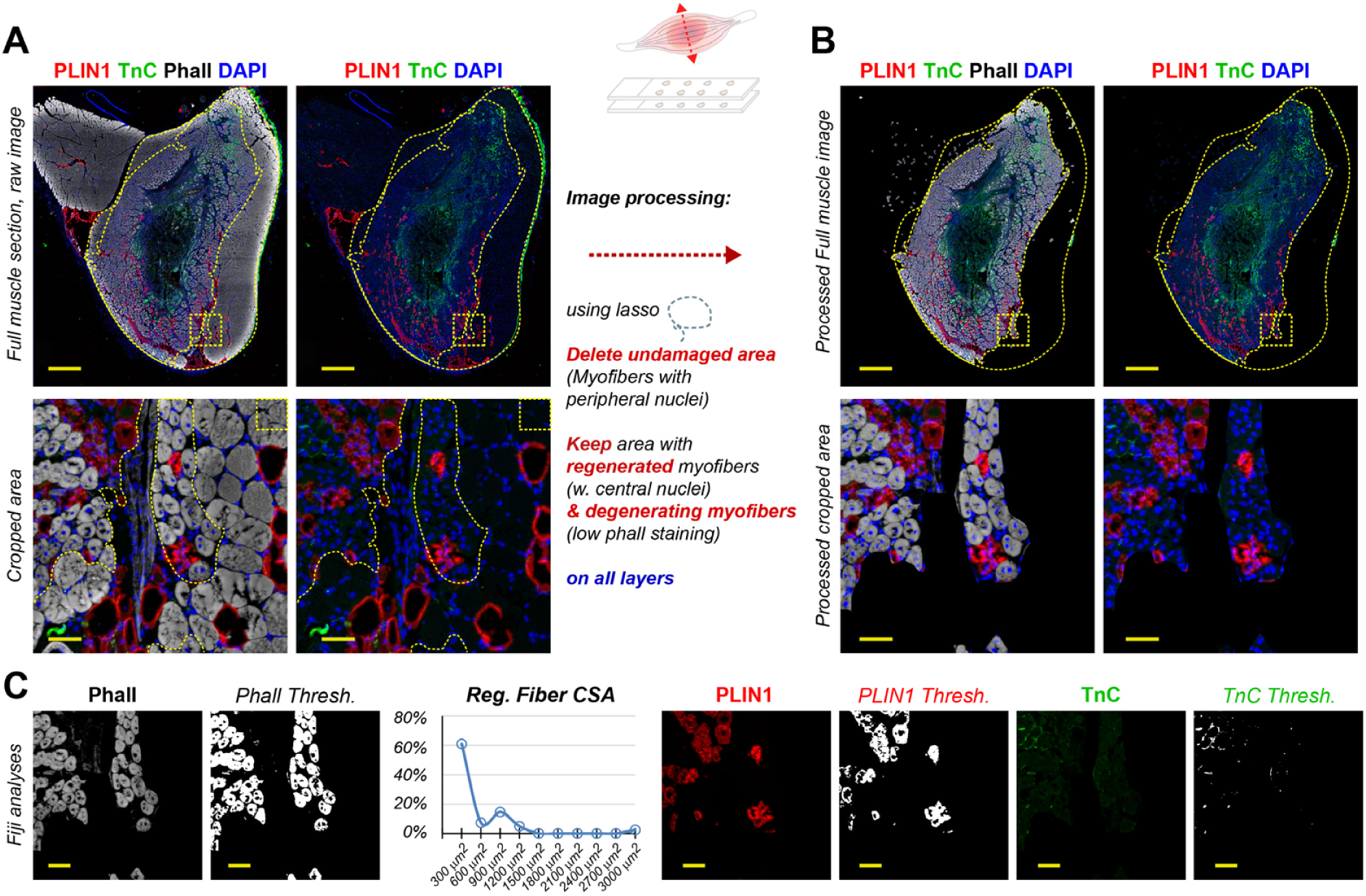
Image analysis pipeline to explore muscle regeneration and IMAT formation in muscle samples after glycerol injury. Muscles collected from injured mice are embedded and serially sectioned, and immunostained with antibodies for morphometric analyses (the examples shown here were stained with phalloidin (white), perilipin1 (PLIN1, red), TnC (green), and with DAPI (blue). The example shown here shows a muscle cross-section from a control mouse, 7 days after glycerol injury, to illustrate the image analysis pipeline used for the following quantifications. (A) Unprocessed images of the whole transverse muscle section (top), or high magnification (bottom) of an area indicates with the dotted square in top images), with (left) or without (right) the phalloidin staining, highlighting with dotted lines the full muscle area analyzed, and the lesion limits as deduced from the transition between unaffected myofibers and regenerated myofibers. (B) Image processing steps applied to each muscle section in order to restrict morphometric quantifications to the lesion only. (C) Images of the same muscle section as in (A), after the image processing steps described in (B), with (left) or without (right) the phalloidin staining, showing the whole muscle section (top) and high magnification images (bottom). (D) view of the same area indicated by dotted square, separating the color channels and showing raw colours or thresholded images (Fiji or ImageJ) from which quantifications are made, as well as an example of plot of the distribution of fiber cross-section areas corresponding to the phalloidin staining in this small image. Scalebars: (A,B, top images) 500 µm; (A, B, bottom images) 50 µm; (C) 50 µm.

Inflammatory invasion and myogenic regeneration start from the outer limits of the lesion, and progress towards the center, such that in a same muscle/stage, regenerated myofibers at the periphery are the oldest and largest, whereas younger regenerated myofibers present in more central areas are smaller (Figure 1E, Figure S1). A ring of active regeneration occurs at the transition between the outer area containing regenerated myofibers, and the central area with necrotic/degenerating myofibers (Figure 1E-F, Figure S1, Figure 3). As regeneration proceeds, this active zone progresses by moving towards the center of the lesion, such that the necrotic area progressively shrinks, while the lesion area expands and is progressively filled with *de novo* regenerated myofibers, which also expand in diameter until completion of the process. In healthy mice, while the necrotic area is still present at 7dpi, it has largely disappeared at 14dpi (Figure 4C,D).

**Figure 4:**
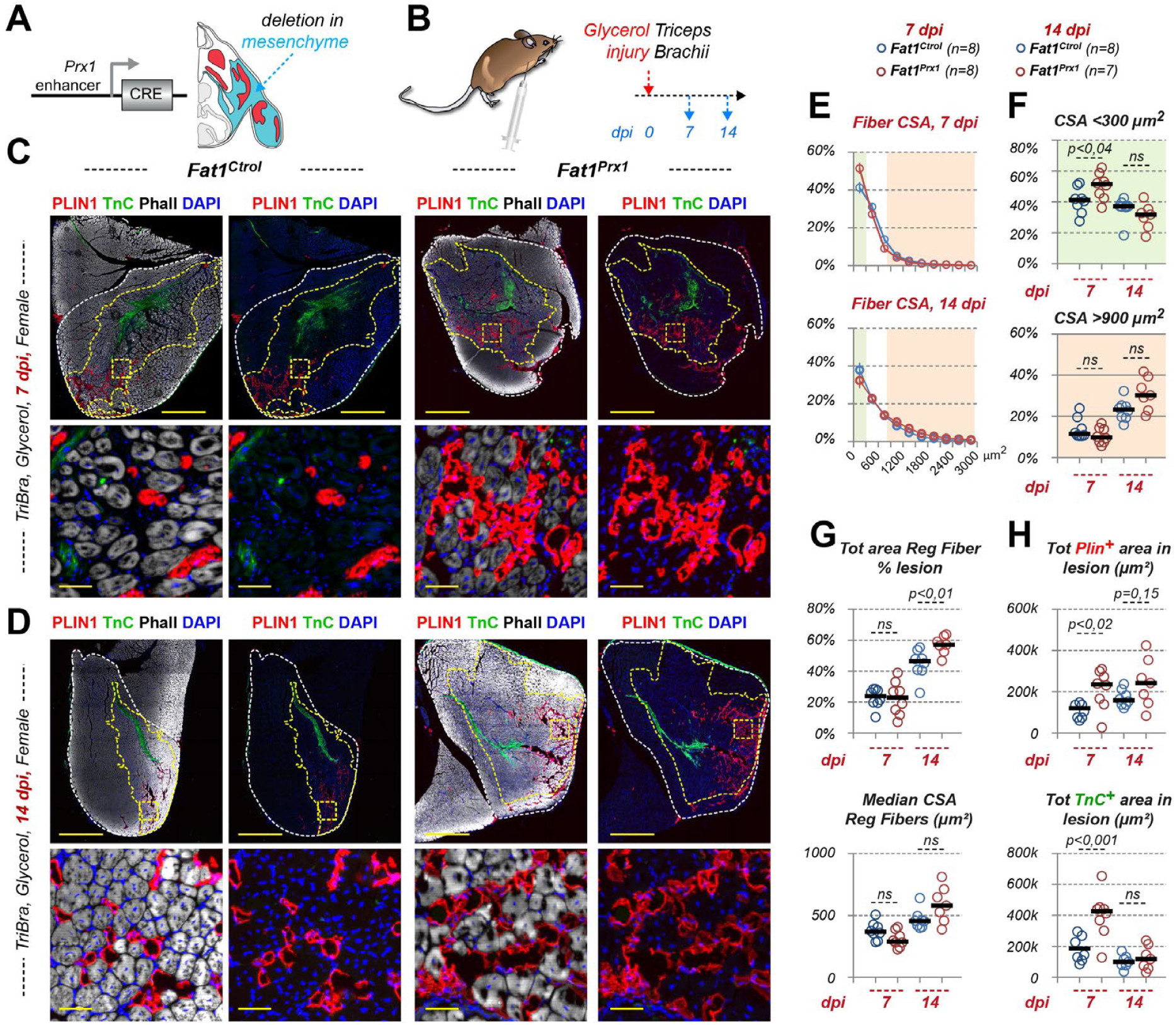
Mesenchyme-specific *Fat1*-deletion enhances glycerol-induced fibro-adipose infiltrations while preserving muscle regeneration. (A) Scheme of the *Prx1-cre* driver used for recombination. The domain of activity of *Prx1-cre* at embryonic stages includes mesenchyme derived from the lateral plate mesoderm, represented in blue, whereas muscles, derived from paraxial mesoderm (somites) are shown in red (the scheme is based on previously published data in (Helmbacher, 2018)). (B) Experimental procedure and timeline: glycerol injury was performed in 4-8 months old mice, and muscles were harvested at 7 or 14 dpi, cryoembedded and serial sectioned. (C, D) Muscle sections were immunostained with phalloidin (white) and antibodies against Perilipin (PLIN1, red) and against Tenascin C (TnC, green) to visualize muscle fibers, adipocytes and transient fibrosis, respectively, in *Fat1^Ctrol^* and *Fat1^Prx1^* female mice, 7 days (C), and 14 days (D) after Glycerol-induced damage in the triceps brachii. Yellow dotted lines highlight the outer limit of the triceps brachii muscle and the limit of the lesion area, while the squared-dotted areas highlight the areas shown at higher magnification in panels below the full muscle views. For each genotype, images are shown with (left) or without (right) the phalloidin staining. The images shown are representative of the female phenotype detailed in Figure 5A-D. (E) Distribution of myofiber cross section areas (CSA), showing ranges up to 3000 µm^2^ (higher CSA ranges, above 3000 µm^2^, were skipped because of low percentages at these stages). (F) Proportions of myofibers in the 0 to 300 µm^2^ range (upper plot, highlighted in green in C), and in the 900 to 2500 µm^2^ range (lower plot, highlighted in light orange in C), showing individual data points for each mouse. (G) Quantifications of the percentage of the lesion area covered by regenerated fibers (top) or median fiber Cross-section-area, showing individual data points for both genotypes and stages. (H) Quantifications of the total area in the lesion covered by PLIN1 staining (top), or by TnC staining (bottom), showing individual data points for both genotypes and stages. Scalebars: (C,D, top images) 1000 µm; (C,D, bottom images) 50 µm.

Given this positional evolution (large fibers in the periphery and small towards the active zone) and the variability of the initial lesion size, we established a systematic procedure for unbiased image quantitative analyses of the phenotypes of our samples. This involves acquiring a mosaic image of the entire muscle cross section, and systematically including phalloidin staining in one color channel. This strategy enabled us to unambiguously delineate the limits of the lesion area, identified by the clear boundary between undamaged fibers with peripheral nuclei and regenerated fibers featuring central (internal) nuclei (Figure 3). The region containing undamaged fibers was manually excluded from all channels including phalloidin but also channels with other antibody signals, so that subsequent ImageJ/Fiji analyses were restricted solely to the lesion area (Figure 3B). This approach allowed us to perform morphometric analyses accross the entire muscle lesion, thereby avoiding the selection bias inherent to analyzing smaller, arbitrarily chosen, muscle areas. This method was used to quantify myofiber cross section area, as well as additional markers used, such as Perilipin-1 (PLIN1) to assess adipogenic infiltrations and Tenascin-C (TnC) as a marker of pro-remodelling FAPs, characteristic of transient fibrosis. In all cases, measurements were compared between genotypes, either as raw value or normalized to the muscle or to the lesion area.

To investigate the impact of *Fat1* deletion in the mesenchymal lineage, we used first the *Prx1-cre* line, which drives recombination throughout the lateral plate-derived mesenchymal lineage from embryonic stages onward (Logan *et al*, 2002). Although *Prx1-cre; Fat1^Flox/Flox^* mice (referred to as *Fat1^Prx1^* mice hereafter) display developmental phenotypes affecting the shape and size of several muscles (Helmbacher, 2018), the TB muscle is relatively unaffected. Our cohort includes mice aged 4 to 9 months (with no difference in age distribution between controls and mutants). This enabled us to assess damage-induced regeneration, in a context with minimal pre-injury phenotypes.

To evaluate muscle regeneration following *Fat1* ablation in the *Prx1-cre* lineage, we measured the cross-section area (CSA) of the regenerated fibers (Figure 4E,F), and the total area occupied by regenerated myofibers within the lesion (Figure 4G). Unlike the robust non-cell autonomous impact of mesenchymal *Fat1* ablation on developmental muscle growth during development (Helmbacher, 2018), the impact on adult regeneration was modest. At 7 dpi, *Fat1^Prx1^* mice exhibited a small increase in the proportion of Fibers with the smallest caliber (<300µm^2^) compared to controls (Figure 4E-G). This did not impact the overall median fiber CSA (Figure 4G top), and was a transient effect not observed at 14dpi. Furthermore, at 14dpi, *Fat1^Prx1^* mice exhibited a subtle but significant increase in the percentage of lesion area covered by regenerated fibers relative to controls (Figure 4G bottom). These regeneration-associated changes are distinct from pre-existing (pre-injury) phenotypes, which manifested as a reduction of the whole muscle section area in *Fat1^Prx1^* mice compared to controls at 7 but no longer at 14 dpi (Figure S3A-C), whereas the size of the lesion (measured as % of the whole muscle area) was not significantly affected by the genotype.

While myogenic repair was largely preserved, quantifications of PLIN1 and TnC localization and abundance uncovered more pronounced effects on FAP-derived fibro-adipogenic cell types: *Fat1^Prx1^* mice exhibited enhanced adipose infiltrations compared to controls (Figure 4H), evident both at 7 dpi, and 14 dpi, and a substantial increase in the TnC+ area at 7dpi. Consistent with the fact that TnC labels a subpopulation of FAPs characteristic of early post lesion stages, this increase in mutants was only observed a 7 dpi and was no longer present at 14dpi, a stage at which the only residual TnC staining was restricted to intramuscular tendon structures (Figure 4D), also present in uninjured muscles (not shown). Further analyses carried out separately on males and female cohorts uncovered that the effect of *Fat1^Prx1^* ablation on adipose infiltrations were more pronounced in females than in males (Figure 5A), whereas the increase in TnC+ area only reached statistical significance in male mutants (Figure 5B,E). The subtle and transient reduction of fiber CSA was likewise only significant in males (Figure 5D), whereas the proportion of the lesion occupied by regenerated fibers at 14 dpi was increased only in females (Figure 5C). These observations indicate that these phenotypes are independent of each-other. Interestingly, the latter increase in proportion of the lesion area covered by regenerated fibers in females co-occurred with an increase in adipocyte-covered area, indicating that in the present study, enhanced adipogenesis was not inversely correlated with regeneration as previously described (Norris *et al*., 2025).

**Figure 5:**
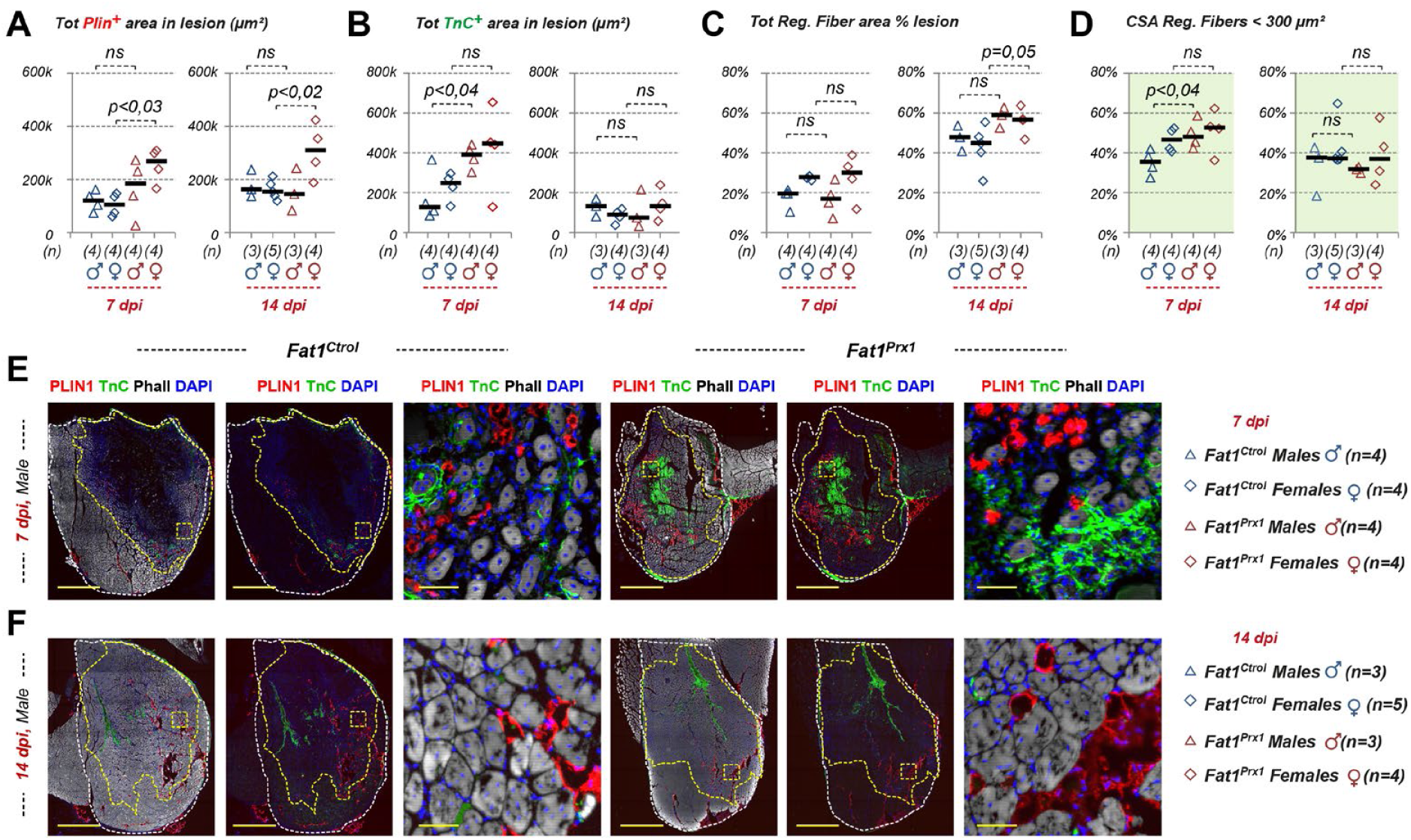
Sex-specific bias in phenotypes induced by mesenchyme-specific Fat1 ablation in muscle after glycerol injury. (A-D) The same quantifications as in Figure 4, of (A) the total PLIN1^+^ area in the lesion, (B) the Total TnC^+^ area in the lesion, (C) the percentage of the lesion area covered by regenerated myofibres, and (D) the percentage of small caliber myofibers (<300µm^2^), showing individual data points for each mouse, at 7 and 14 dpi, and separating males and females (genotypes and number of mice in each are shown at the bottom right of the figure. (E) Sections of *Fat1^Ctrol^* and *Fat1^Prx1^* male mice, 7 days (C), and 14 days (D) after Glycerol-induced damage in the triceps brachii were immunostained with phalloidin (white) and antibodies against Perilipin (PLIN1, red) and against Tenascin C (TnC, green) to visualize muscle fibers, adipocytes and transient fibrosis. The images shown are representative of the male phenotype detailed in (A-D). Scalebars: (E,F, for each genotype, the first two images) 1000 µm; (E,F, for each genotype, right side image) 50 µm.

Overall, our results show that mesenchymal *Fat1* activity is largely dispensable for the promyogenic activity of FAPs during skeletal muscle repair, as its deletion minimally impacts the efficacy of muscle regeneration. Instead, *Fat1* activity in the mesenchymal lineage is required to limit the expansion of fibro-adipose infiltrates triggered by acute glycerol injury. Our analyses also uncovered sex-specific aspects of the phenotypes resulting from mesenchymal *Fat1* depletion, with a female-bias for the enhanced adipose infiltrations and the slightly enhanced regenerated area, and a male bias for the enhanced TnC expansion and for the transient effect on fiber CSA. These observations also imply that at the stages studied, the *Fat1*-dependent enhancement of adipose infiltrations is not sufficient to interfere with the efficacy of regeneration.

### Cell-autonomous and cell-non-autonomous effects of inducible Fat1 deletion in FAPs

To further understand the role played by *Fat1* in the formation and expansion of fibro-fatty infiltrates, and completely suppress the confounding pre-injury phenotypes, we used a second CRE line, referred to as *Pdgfra^icre^*, in which expression of the Tamoxifen-inducible Cre/ERT2 (iCRE) is driven by exogenous *Pdgfra* regulatory genomic locus in a BAC Transgene, allowing inducible deletion in the *Pdgfra* lineage (Rivers *et al*, 2008). Although *Pdgfra*-positive mesenchymal cells are also present outside muscles, *Pdgfra* expression in muscle encompasses all FAP subtypes and is the most widely used marker of FAPs used for FACS studies. A cohort of adult control (*Fat1^Ctrol^; Pdgfra^icre^; R26^YFP/+^*, referred to as *Ctrol^iPdgfra-YFP^*) and mutant (*Fat1^Flox/Flox^; Pdgfra^icre^; R26^YFP/+^*, referred to as *Fat1^iPdgfra-YFP^*) mice (Figure 6A), was subjected to a regime of 5 consecutive Tamoxifen injections (1 per day), starting 2 days prior to Glycerol muscle lesions in the triceps brachii muscle at day 0 (Figure 6B). Analyses were carried out at 7 and 14 dpi.

**Figure 6:**
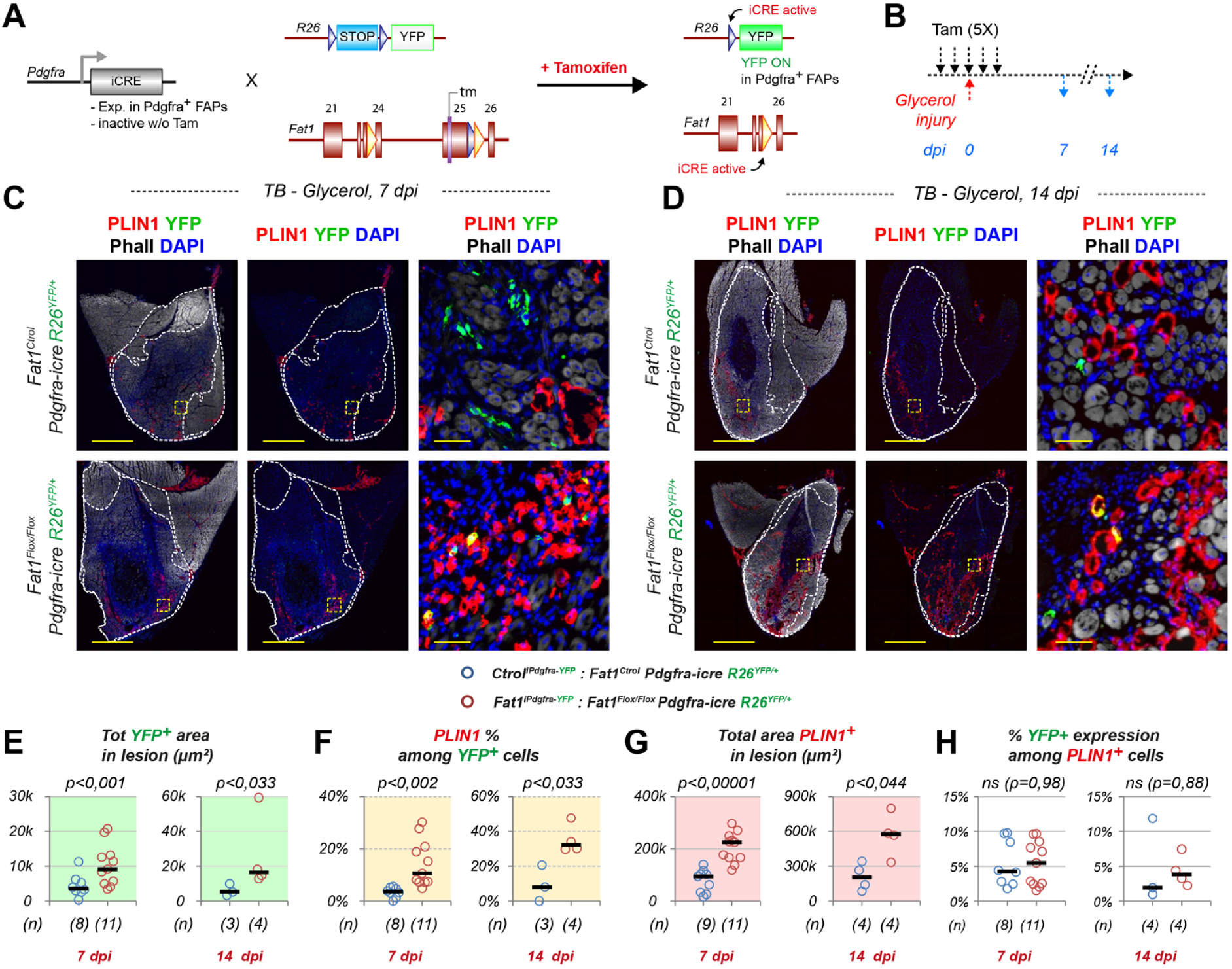
inducible *Fat1*-deletion in the Pdgfra lineage leads to cell-autonomous and non-cell-autonomous enhancement of adipogenic differentiation in glycerol-injured muscles. (A) Genetic paradigm used for inducible-recombination in the Pdgfra lineage, with Pdgfra-iCRE line, combined with the Fat1-Flox and Reporter allele R26-YFP. Upon addition of Tamoxifen, activated CRE acts on both alleles, and YFP expression is used to follow cells in which recombination occurred. (B) Scheme of the experimental design, with d0 representing the onset of experimentation, on adult mice of the indicated genotypes. Tamoxifen is applied 5 consecutive days, starting 2 days before muscle damage. Analysis is performed 7 or 14 days post-damage. (C, D) Sections of damaged triceps brachii muscles collected from control (*Fat1^control^; Pdgfra -icre; R26^YFP/+^; abbreviated as Ctrol^iPdgfra-YFP^*, top images) and FAP-specific *Fat1* mutant mice (*Fat1^Flox/Flox^; Pdgfra -icre; R26^YFP/+^*, *abbreviated as Fat1^iPdgfra-YFP^*, bottom images), at 7dpi (C) or 14 dpi (D). Sections were immunostained with antibodies to GFP/YFP (green), Perilipin1 (PLIN1, red), with Fluorescent-Phalloidin (white) and DAPI (blue). For each genotype/stage, the two left panels show an overview of the entire muscle cross-section, while the right panel show a higher magnification of the area highlighted with squared dotted lines. For each muscle, we highlighted with dotted lines the outer limit of the studied muscle (triceps brachii, median), and the lesion area (defined by the presence of myofibers with central nuclei (indicating regeneration). (E, F, G, H) Quantifications of the total YFP+ area (E) as a proxi for the number of *Pdgfra-icre*-derived YFP^+^ cells, of the PLIN1^+^ percentage among the YFP+ area (F), the total PLIN1+ area (G), and the YFP^+^ percentage among PLIN1^+^ area (H), in the lesion, comparing *Ctrol^iPdgfra-YFP^*, and *Fat1^iPdgfra-YFP^*mice. Each dot represents and individual mouse, and the number of mice is indicated by the numbers between brackets below the plots. Scalebars: (C,D, for each genotype, the first two images (left and center)) 1000 µm; (C, D, for each genotype, right side image) 50 µm.

In these mice, the *R26^YFP^* reporter line was used as a readout of recombination, allowing to visualize YFP-positive cells in the interstitial space between muscle fibers, labelling FAPs and their differentiation progeny (Figures 6 and 7), and to compare the fate of recombined cells in control and mutant contexts. Most YFP^+^ cells were concentrated in the lesion, while largely absent from unaffected muscle areas, likely because 3 of the 5 Tamoxifen injections were done after glycerol injection, which triggers FAP expansion, thereby mobilizing the *Pdgfra* locus in the lesion. This regime resulted in low recombination efficacy, with few YFP^+^ cells per section, much lower than the recombination rate observed with the same *Pdgfra^icre^* line at embryonic and post-natal studies (Helmbacher, 2018, 2022), and also lower than the recombination rate described for *Pdgfra^icre^* lines used in other FAPs studies (Kajabadi *et al*, 2023; Norris *et al*, 2023). We had excluded working with these alternative lines, because they involved knockout alleles of the endogenous *Pdgfra* locus, raising concerns that *Pdgfra* heterozygosity could influence or exacerbate phenotypes related to the pathway analyzed. Importantly, the low numbers of *Pdgfra*-derived YFP^+^ cells per section obtained in our study offered us the advantage to allow distinguishing cell autonomous from non-cell autonomous phenotypes.

**Figure 7:**
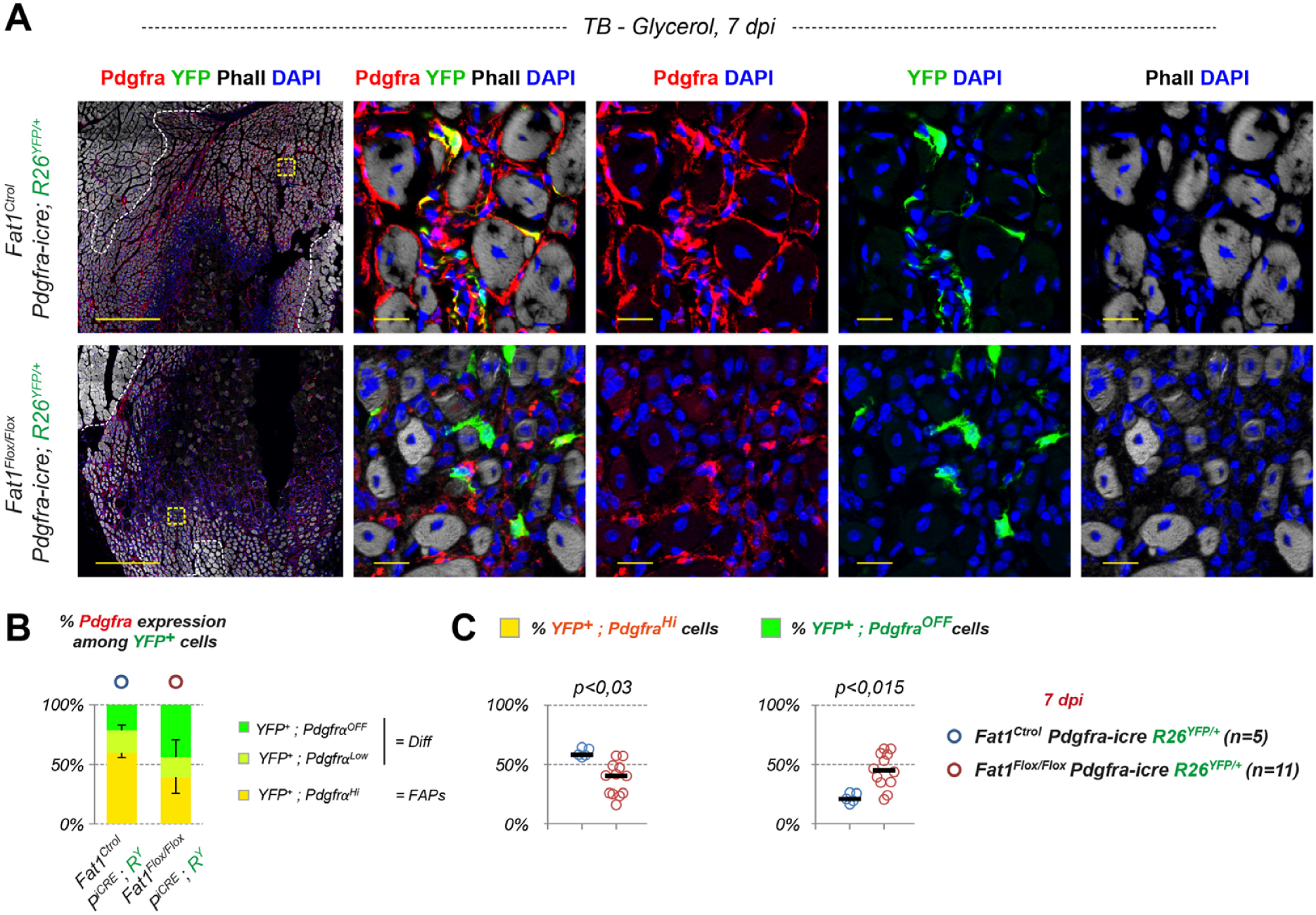
inducible *Fat1*-deletion in the Pdgfra lineage leads to expansion of Pdgfra-derived lineage and loss of Pdgfra expression. **(A)** Sections of damaged triceps brachii muscles from control (*Fat1^control^; Pdgfra -icre; R26^YFP/+^*, top images) and FAP-specific *Fat1* mutant mice (*Fat1^Flox/Flox^; Pdgfra -icre; R26^YFP/+^*, bottom images), immunostained with Phalloidin (white), DAPI (blue), and antibodies to YFP (green), and Pdgfra (red). The yellow dotted lines demarcate the lesion area from unaffected muscle tissue. White squares indicate the position of higher magnification images shown in Figure 4. **(D)** Quantification of percentages among YFP^+^ cells, that express high levels (yellow), low levels (light green) or no Pdgfra (dark green, Pdgfra^OFF^) in *Ctrol ^iPdgfra-YFP^* and *Fat1^iPdgfra-YFP^* mice. **(D)** Plots of the Pdgfra^High^ percentage (left), and Pdgfra^OFF^ percentage (right) among the YFP+ area, showing individual data points for each mice. Scalebars: (A, overview) 500 µm; (A, crops) 20 µm.

As previously observed in our work on migrating retinal astrocytes (Helmbacher, 2022), inducible deletion of *Fat1* in the *Pdgfra* lineage led to an increased the number of YFP^+^ cells in *Fat1^iPdgfra^* mice compared to *Ctrol^iPdgfra^* mice (Figure 6E), consistent with an effect on cell proliferation. Given the enhanced adipogenesis phenotype observed in the *Prx1-cre* model, we next analyzed the adipogenic fate of recombined cells in the inducible *Pdgfra* model, combining anti-GFP/YFP with anti-perilipin1/PLIN1 antibodies, and quantified the ratio of PLIN^+^;YFP^+^ versus total YFP^+^ area. This uncovered a significant increase in adipogenic fate of recombined cells in *Fat1^iPdgfra-YFP^* mice compared to *Ctrol^iPdgfra-YFP^* mice (Figure 6C,D,F). The severity of this phenotype was variable, with the percentage of PLIN^+^;YFP^+^ area ranging from 5% to 30% (median 10%) in *Fat1^iPdgfra^* mice at 7 dpi, while *Ctrol^iPdgfra^* mice ranged between 0 and 6% (median 3%). At 14 dpi, these values increased from 30% to 48% in *Fat1^iPdgfra^* mice (median 32%), while *Ctrol^iPdgfra^* mice ranged between 0 and 21% (median 7,8%).

This increase in the propensity of recombined cells to undergoe adipogenic differentiation in *Fat1^iPdgfra-YFP^* mice was accompanied with a decrease in the proportion of YFP^+^ cells maintaining PDGFRA protein expression (Figure 7). The loss of Pdgfra expression likely reflects the loss of progenitor characteristics and is consistent with the fact that mature adipocytes do not maintain *Pdgfra* expression (Contreras *et al*, 2019). The proportion of PDGFRA-negative; YFP^+^ cells in *Fat1^iPdgfra-YFP^* mice was higher than the proportion of PLIN^+^; YFP^+^ cells, suggesting that *Fat1* deletion may also enhance alternative FAP differentiation outcomes, such as myofibroblasts.

Unexpectedly, the analysis of PLIN1/YFP double stainings also uncovered a robust non-cell-autonomous enhancement of adipogenesis: The total PLIN1^+^ area in the lesion, consisting of a large majority of non-recombined cells (YFP-negative), was also significantly increased (Figure 6G). Although it is theoretically possible that recombination at the *R26* and *Fat1* loci may not systematically occur in the same cells, if this increase in adipocyte number resulted from the cell autonomous phenotype described above, we would nevertheless expect an increase in the YFP+ proportion of the total PLIN+ area in mutants. Instead, the proportion of YFP^+^ among PLIN1^+^ cells remained constant, at around 5% in both control and mutant mice (Figure 6H), supporting the idea that *Fat1*-deficient FAPs exert a pro-adipogenic influence on neighbouring non-recombined FAPs, or have lost an inhibitory (anti-adipogenic) property.

In line with our observations in the *Fat1^Prx1^* model, analysis by sex revealed that enhanced adipogenesis was more pronounced in female *Fat1^iPdgfra-YFP^* mice compared to *Ctrol^iPdgfra-YFP^* females than in males (Figure 8A middle graph). The effect size was approximately 5 folds in females, versus 2.3 folds in males. In contrast, the non-cell autonomous increase in adipogenesis was similar (around 2,5 folds) between males and female (Figure 8A right graph), supporting the idea that the two phenotypes are distinct from each other and not equally sensitive to sex.

**Figure 8:**
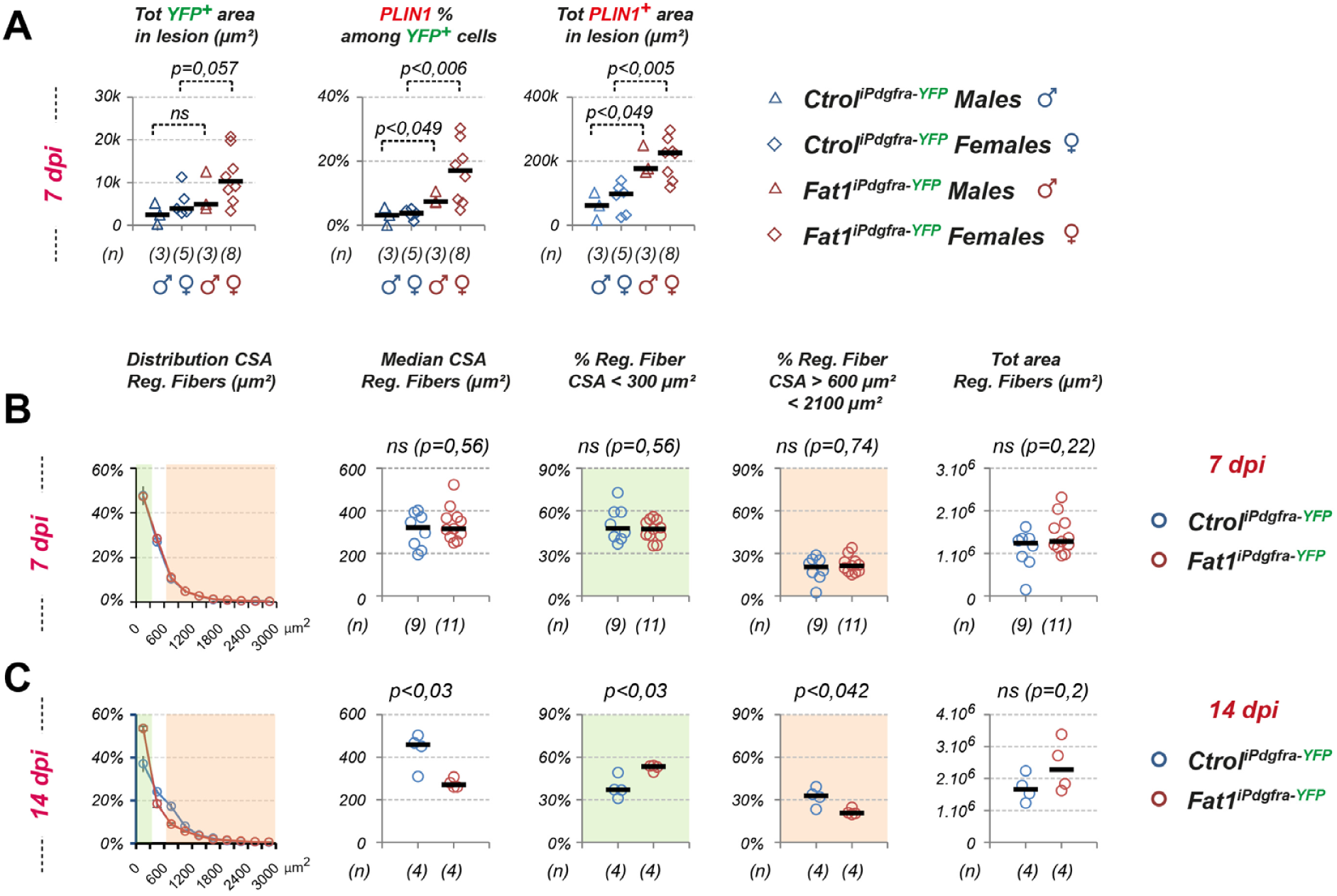
Female-specific enhancement of intramuscular adipogenesis and effects on myogenic repair in the inducible *Pdgfra-icre* model. (A) The same quantifications as in Figure 5E-G, of the total YFP^+^ area in the lesion (Left plot), the PLIN1^+^ proportion of the YFP^+^ area (middle plot), and the total PLIN1^+^ area in the lesion (right plot), showing individual data points for each mouse, and separating males and females (genotypes and number of mice area shown below the graph), comparing *Ctrol^iPdgfra-YFP^* and *Fat1^iPdgfra-YFP^* male and female mice, at 7 dpi. (B, C) Quantifications of morphometric parameters related to regeneration in *Ctrol^iPdgfra-YFP^*and *Fat1^iPdgfra-YFP^* mice, at 7 dpi (B) and 14 dpi (C), including from left to right: 1) the distribution of myofiber cross section areas (CSA), showing ranges up to 3000 µm^2^, 2) the median CSA, 3) the proportions of myofibers in the 0 to 300 µm^2^ range (highlighted in green in the left plots), 4) the proportion of myofibers in the 600 to 2100 µm^2^ range (highlighted in light orange in the left plots), 5) the total area covered by regenerated fibers in the lesion, showing individual data points for both genotypes and stages.

Finally, given the strong non-cell-autonomous effect on adipogenesis, despite modest recombination rates, we next assessed whether inducible *Fat1* deletion in FAPs impacts muscle regeneration by myofiber CSA and total regenerated area. We did not detect any effect on fiber CSA or total area covered by regenerated fibers at 7dpi, contrasting with the changes observed in the *Fat1^Prx1^* model. Instead, at 14 dpi, this analysis uncovered a significant reduction of the median fiber CSA in *Fat1^iPdgfra-YFP^* mice compared to *Ctrol^iPdgfra-YFP^* mice, resulting not only from an increased proportion of fibers with small caliber (<300µm^2^), but also from a decreased proportion of large diameters (>600µm^2^; <2100µm^2^) (Figure 8B). Although we cannot distinguish whether this effect is caused by the enhanced adipogenesis, or by the loss of a promyogenic *Fat1* activity in FAPs, this observation suggests that inducible *Fat1* deletion in FAPs negatively interferes with myogenic repair, an effect that was possibly masked in the *Fat1^Prx1^* model by pre-injury (developmental) phenotypes (which are absent in the inducible *Fat1^iPdgfra-YFP^* model (Figure S3), as Tamoxifen-induction is coincident with the lesion).

## Discussion

The appearance of intramuscular fibrosis and adipose tissue is a hallmark of advanced symptoms in muscles of patients with muscular dystrophy (Flores-Opazo *et al*., 2024; Theret *et al*., 2021). Although common to multiple diseases, the appearance of fibro-fatty infiltrates is frequently a consequence of chronic inflammation, itself being a secondary to the primary cause of each disease. So far, to our knowledge, conditions in which FAP dysfunction is the primary event have not been identified in human patients. Therefore, animal models engineered to primarily affect FAP homeostasis represent alternative sources of key information to understand what contributes to fibro-fatty development and to identify novel therapeutic approaches, potentially applicable to multiple disease conditions. While FAP-specific deletion of *Fat1* was initially aimed to explore the impact on the efficiency of muscle regeneration, we made the unexpected discovery that it enhanced fibro-adipogenic differentiation in regenerating muscles after glycerol injury, with a strong bias towards adipogenesis, including both a cell autonomous and a non-cell-autonomous component. This uncovered a new role of *Fat1* as an inhibitor of intramuscular fibro-adipogenesis, and positions *Fat1^Prx1^* mice as a new model of enhanced IMAT formation in response to injury.

### Distinguishing developmental from regenerative promyogenic activities of FAPs

FAPs constitute a promyogenic niche that supports regeneration of adult skeletal muscles by secreting factors promoting satellite cell expansion or myogenic repair (Joe *et al*., 2010; Lukjanenko *et al*., 2019; Madaro *et al*., 2018), by secreting chemokines orchestrating key steps in the FAP-immune cross-talk (Heredia *et al*., 2013; Lemos *et al*., 2015; Nawaz *et al*, 2022), and by structuring the extracellular matrix, influencing myogenesis through its composition and its stiffness (Kotsaris *et al*, 2023). Likewise, during embryonic development, a population of muscle-associated mesenchymal cells plays similar promyogenic functions (Helmbacher & Stricker, 2020) and was shown to represent the developmental progenitors of adult FAPs (Vallecillo-Garcia *et al*., 2017), the two lineages sharing expression of the transcription factor Osr1 (Kotsaris *et al*., 2023; Stumm *et al*., 2018; Vallecillo-Garcia *et al*., 2017). Genetic ablation of adult FAPs resulted in reduced myogenic stem cell expansion after muscle injury (Murphy *et al*, 2011; Wosczyna *et al*., 2019), and consequently in muscle atrophy and weakness, or in impaired revascularization after hindlimb ischemia (Santini *et al*., 2020). Similarly, abrogating fate or functions of developmental FAPs has an influence on muscle growth and patterning, illustrated in multiple mesenchyme-specific mouse mutants (Helmbacher & Stricker, 2020). Conditioned media from adult FAPs extracted from injured muscles release factors that promote myogenesis (Joe *et al*., 2010; Madaro *et al*., 2018; Mozzetta *et al*., 2013). While FAP-derived secretomes have been explored in several studies (Florin *et al*, 2020; Kotsaris *et al*., 2023; Vumbaca *et al*, 2021), only a small fraction of these promyogenic factors have so far been functionally identified and confirmed to promote muscle stem cell growth or myofiber growth. This is the case for WISP1, BMP3a/GDF10, or Follistatin, three factors produced by FAPs in young but not old mice, which decreased production in aged dystrophic mice is responsible for myofiber atrophy and muscle weakness (Lukjanenko *et al*., 2019; Nawaz *et al*., 2022; Uezumi *et al*., 2021). Similarly, exploring the promyogenic transcriptome or secretome of developmental FAPs has yielded a number of potential mediators of their promyogenic activity (Besse *et al*, 2020; Orgeur *et al*, 2018; Vallecillo-Garcia *et al*., 2017).

During development, abolishing mesenchymal *Fat1* activity had a strong effect of muscle development, impacting progenitor migration and fiber extension in the cutaneous maximus (CM) muscle, but also leading to the presence of ectopic muscles with unconventional fiber orientation and attachment sites (Helmbacher, 2018). These phenotypes recapitulated a large subset of the muscle phenotypes observed in the constitutive knockout (Caruso *et al*., 2013). We thus anticipated that abolishing mesenchymal *Fat1* in adult muscles would likewise interfere with the promyogenic activity of FAPs. However, mesenchymal *Fat1* ablation (in both constitutive and inducible models) only had a relatively modest (even though significant) effect on myogenic repair, limited to a small increase in the proportion of small caliber fibers, and to the reduced myofiber CSA observed at 14 dpi in the inducible *Fat1^iPdgfra^* model. Although at fetal stages, *Fat1* levels were higher in the TB muscle than in other developing muscles (Mariot *et al*., 2015), it remains possible that in adult mice, such a promyogenic activity might vary between muscle types, and would be relatively mild in the triceps brachii, while more pronounced in other muscles (that we have not sought to identify). An alternative possibility could be that the *Fat1*-regulated mesenchymal factors influencing muscle morphogenesis during development might be distinct from the FAP-derived pro-myogenic factors that promote myogenic repair in adults. Considering that ablation of mesenchymal *Fat1* resulted instead in a robust increase in the amount of intramuscular adipose differentiation, resulting from the combination of cell-autonomous and non-cell autonomous component, this suggests that *Fat1* activity in adult FAPs may have been redirected toward inhibiting adipogenesis, at the expense of a role in promoting secretion of promyogenic factors. It is also possible that the nature of *Fat1*-regulated FAP transcriptome (or secretome) might depend on the molecular properties of FAPs, and that the FAP subtypes present in TB muscle after glycerol injury might be distinct from embryonic FAPs surrounding *Fat1*-dependent muscles during development. In support of this possibility, it is interesting to consider *Osr1*, another example of genes acting in both embryonic and adult FAPs (Kotsaris *et al*., 2023; Stumm *et al*., 2018; Vallecillo-Garcia *et al*., 2017). Unlike *Fat1* however, *Osr1* deletion had an impact on both developmental and regenerative myogenesis (Kotsaris *et al*., 2023; Vallecillo-Garcia *et al*., 2017). Nevertheless, as postulated above, the *Osr1*-dependent transcriptome signatures appear qualitatively different between developmental and adult stages (Kotsaris *et al*., 2023; Vallecillo-Garcia *et al*., 2017). Consistently, the single cell transcriptome studies that have characterized molecularly distinct FAPs subpopulations, representing phases of regeneration and modalities of FAP activities, differently represented in healthy or pathological environments, have also identified differences between ages, accounting for differences between young and aged FAPs, but also differences between juvenile and adult FAPs.

### Fat1 is a negative regulator of intramuscular adipogenesis

The present findings identify *Fat1* as a new modulator of fibro-adipogenic differentiation. This adds to the recent identification of a series of other signaling pathways that significantly moderate adipogenic FAP differentiation, among which interleukin-4 (IL4) (Dong *et al*, 2014) Nitric oxid (NO) and Notch signaling (Marinkovic *et al*, 2019), Desert Hedgehog (DHH) (Kopinke *et al*, 2017; Norris *et al*., 2023), Wnt5A, and Wnt7A (Fu *et al*, 2023; Santiago *et al*, 2025). After muscle injury, FAPs transiently harbor cilia, which represent a hotspot of active Hedgehog signaling and are necessary for intramuscular adipogenesis (Kopinke *et al*., 2017). Genetically preventing cilia formation in FAPs, deleting Patched1 activity in FAPs, or activating Hedgehog signaling with the Smoothened agonist SAG, equally resulted in a reduction of intramuscular adipogenesis, and led to increased diameter of regenerated fibers (Kopinke *et al*., 2017). A follow-up study identified Desert hedgehog as the ligand, produced by endothelial cells and schwann cells, which limits adipogenesis in regenerating muscle (Norris *et al*., 2023). In another study, an in vitro screen using FAPs from dystrophic mice led to the finding that inhibitors of GSK3 were overrepresented among anti-adipogenic compounds, suggesting that canonical Wnt signaling could inhibit intramuscular adipogenesis (Reggio *et al*, 2020). ScRNAseq analyses indicated that FAPs were the main source of Wnt ligands. Among them, WNT5a was expressed in resting and lesion-activated FAPs, downregulated in MDX FAPs, and capable of inhibiting adipogenesis of FAPs in vitro, (Reggio *et al*., 2020). Another FAP-derived Wnt, WNT7A, previously known to promote muscle regeneration induced by injury by inducing expansion of satellite cells (Le Grand *et al*, 2009), and to improve regeneration in dystrophic mice (von Maltzahn *et al*, 2012), while its ablation exacerbated muscular dystrophy symptoms (Gurriaran-Rodriguez *et al*, 2024), was recently shown to suppress intramuscular adipogenesis in vitro and in vivo (Fu *et al*., 2023). Furthermore, injection of lipid nanoparticles loaded with WNT7A-mRNA in mice were shown to be effective in reducing IMAT in vivo (Santiago *et al*., 2025).

In the pathways leading to adipogenesis, adipogenic stem cells are mutipotential, and the first step is a step of commitment, restricting their potential to one fate only (Ferrero *et al*, 2020; Ghaben & Scherer, 2019). This is followed by differentiation and maturation, two subsequent steps that allow maturation of adipocytes into lipid producing cells (Ferrero *et al*., 2020; Ghaben & Scherer, 2019). The master regulator of adipogenic differentiation and maturation is PPARɣ, a nuclear receptor acting as transcription factor. Suppression of PPARɣ activity blocks adipogenic differentiation, and its ablation in all cells except the placenta (Dammone *et al*, 2018) or its inducible ablation in FAPs (Norris *et al*., 2025), abrogate intramuscular adipose tissue formation induced by glycerol lesions. The mechanism by which the anti-adipogenic signals described above act is likely converging on PPARɣ. WNT5A appears to block expression of PPARɣ in a β-catenin-dependent manner (Reggio *et al*., 2020). Interestingly, the anti-adipogenic activity of WNT7A correlates with its capacity to promote nuclear accumulation of YAP/TAZ, both of which acting as anti-adipogenic transcription factors in non-muscular adipose tissue: TAZ acts as a direct suppressor of PPARɣ activity (El Ouarrat *et al*, 2020; Jung *et al*, 2009), whereas YAP may act by inducing WNT5A expression (Lee *et al*, 2022). Paradoxically, loss of *Fat1* activity leads to cell-autonomous nuclear accumulation of YAP and TAZ in skin cancer (Pastushenko *et al*, 2021), and to reduced YAP/TAZ degradation in cancer cells and endothelial cells (Li *et al*, 2023). Thus, based on knowledge mentioned above from non-muscular adipose tissue (El Ouarrat *et al*., 2020; Jung *et al*., 2009; Lee *et al*., 2022), loss-of-*Fat1* would be predicted to cell-autonomously suppress adipogenesis, which is the opposite of what we observed in the present study. However, activities of Fat cadherins, but also YAP/TAZ, vary with parameters such as stiffness changes, position along expression gradients, cell type and activity of neighboring cells, and several arguments support the idea of cell specificity in Fat1 mode of action. First, *FAT1* is both known as a tumor suppressor (Morris *et al*, 2013; Pastushenko *et al*., 2021) or oncogene (Chen *et al*, 2022; de Bock *et al*, 2012; Meng *et al*, 2021; Zhao *et al*, 2025), depending on the cancer type. Second, FAP-derived adipocytes are not identical to adipose-tissue-derived adipocytes, as they differ for example in their insulin sensitivity (Arrighi *et al*, 2015). Third, in the *Fat1^Prx1^* model, we did not observe any sign of obesity (not even a difference in weight), suggesting that the increase in intramuscular adipogenesis after glycerol lesion in *Fat1^Prx1^* mice is not associated with enhanced adipogenesis in non-muscular adipose tissue. Finally, it is interesting to notice that in our tibialis anterior muscle samples, adipogenic infiltrations are regionalized and always found centered around a hotspot of perivascular Fat on the posterior side of the muscle (bottom of full size muscle sections in Figures 3, 4, and 6), while we never found adipocytes in the bone-associated side of the muscle (top of images). This suggest that *Fat1*-restricted adipogenesis is a regionalized process potentially reflecting heterogeneity in FAP populations. Thus, future work will be necessary to characterize the mechanisms by which *Fat1* activity negatively regulates intramuscular adipogenesis and how it may differ from *Fat1*-regulated fibrosis or *Fat1* activity in non-muscular stromal cells.

### Fat1 non-cell-autonomously inhibits intramuscular adipogenesis

An unexpected aspect of our work is the discovery that in addition to cell-autonomoulsy preventing adipogenic differentiation of FAPs, Fat1 also exerts its anti-adipogenic role through a non-cell-autonomous or paracrine activity. This was discovered owing to the limited recombination efficiency of the *Pdgfra-icre* line we used and the Tamoxifen-regime we applied. The fact that we could distinguish adipocytes derived from either recombined cells (YFP+) or non-recombined (YFP-negative) cells, in both controls and mutants, meant that we could differentially quantify cell autonomous and non-cell autonomous phenotypes resulting from *Fat1* ablation. This led to the clear discovery that *Fat1* exerted both types of activities. The non-cell-autonomous phenotype suggests that *Fat1*-deficient FAPs promote adipogenesis of neighboring wild-type FAPs. This effect could result either from an increased secretion of pro-adipogenic signals or from a decrease in secretion of anti-adipogenic signals. Interestingly, while the cell autonomous phenotype was more pronounced in females, the non-cell autonomous part equally affected males and females (Figure 8B). This suggests that the two activities involve distinct molecular mechanisms, not equally sensitive to sex-dependent parameters.

This non-cell autonomous anti-adipogenic paracrine activity is interesting, in the light of studies on adipose tissue, carried out in non-muscular adipose tissue, that identified a subtype of adipogenic stem cells endowed with an anti-adipogenic activity, referred to as Areg (adipogenic regulators), and distinguishable through its expression of F3/CD142, GDF10 (Schwalie *et al*, 2018). The same group later identified several possible mediators of this anti-adipogenic activity, among which GDF10 or Retinoic acid signaling (Zachara *et al*, 2022). Another study later identified a similar subtype of GDF10-expressing FAPs, and found reduced proportions of GDF10+ FAPs in Mdx mice, correlating with enhanced intramuscular fat (Camps *et al*, 2020). These elements suggest that GDF10 exerts a protective role against pathological intramuscular adipose tissue expansion, in addition to its role, mentioned earlier, as FAP-derived signal promoting myofiber growth, and NMJ integrity (Uezumi *et al*., 2021). It will be interesting in future studies to determine how *Fat1* signaling in FAPs is linked to the various players modulating adipogenesis cell-autonomously, and if it indeed modulates secretion of paracrine acting anti-adipogenic factors.

### Sex bias in Fat1-dependent phenotypes

An interesting aspect of our work is that separating male and female cohorts allowed uncovering sex-specific aspects of the *Fat1*-dependent phenotypes. As discussed above, the cell-autonomous phenotype leading to enhanced adipogenicity tends to be associated with females, whereas the increased secretion of the damage-associated molecular pattern TNC is a male-specific response to *Fat1* deficiency. Instead, the non-cell autonomous increase in adipogenesis seen in the inducible model similarly affects both sexes. In contrast, the transient effect on fiber CSA (higher proportion of small caliber fibers) was specifically seen in males, whereas the increased area covered by regenerated fibers at 14dpi was only seen in females. Although modest, these myogenic phenotypes also uncover sensitivity to sex-associated parameters. While there are no reports to our knowledge of sex-specific differences in signaling by Fat-like cadherins, sex-specific associations with IMAT have previously been reported, such as in the remodeling of IMAT in response to muscle injury (Norris *et al*., 2024) or to high fat diet (HFD) (Smith *et al*, 2025).

## Conclusion

Altogether, our results link FAT1 dysfunction in FAPs to IMAT formation, the hallmark of advanced disease progression associated with several pathologies. Although FAT1 itself is not an easily targetable gene for therapy, the discovery of its participation to the gatekeeping system negatively modulating IMAT formation and progression opens new avenues for the search of therapies to block or delay disease progression in patients.

## Methods

### Ethics statement

All animal care and procedures involving mice have been approved by the Ethics committee for animal experimentation of Marseille (committee N° 14, project number 2022011100527909-V2 #34611) and by the French ministry of research and higher education (project number APAFIS #34611-2022011100527909 v3), in accordance with the European Community Council Directives (2010/63/EU) and with the French law and institutional guidelines on animal research. Animal housing, care and experimental protocols were carried out in the IBDM animal facility under an agreement for (G1305521) delivered by the “Prefecture de la region Provence-Alpes-Cotes-d’Azur et des Bouches du Rhônes”.

### Mice

The mouse lines used in this study are maintained with occasional backcrosses with commercial C57BL/6J or B6D2-F1J/RJ mice. We used combinations of the following genetic modifications: *Fat1^Flox^* (*Fat1^tm1.1Fhel^* conditional allele, MGI: 5524120, (Caruso *et al*., 2013; Helmbacher, 2018)); *Prx1-cre* (Transgenic line Tg(Prrx1-cre)1Cjt, MGI: 2450929, (Logan *et al*., 2002)); *Pdgfra-CRE/ERT2* (referred to as *Pdgfra-iCRE*, BAC transgenic line Tg(Pdgfra-cre/ERT2)1Wdr, MGI: 3832569, (Rivers *et al*., 2008)); R26-YFP (reporter line for cre activity, *Gt(ROSA)26sor^tm1(EYFP)Cos^*, MGI: 2449038, (Srinivas *et al*, 2001)); *Fat1^LacZ^* (Bay Genomics Gene trap line *Fat1^Gt(KST249)Byg^*, MGI: 4124012, (Caruso *et al*., 2013; Helmbacher, 2018; Mitchell *et al*, 2001)). To produce Mice for the *Prx1-cre* cohort, *Prx1-cre; Fat1^Flox/+^* males were crossed with *Fat1^Flox/Flox^* females or *Prx1-cre; Fat1^Flox/Flox^* males were crossed with *Fat1^Flox/+^* females. The absence of germline activity of Prx1-cre in males (in contrast to Females) ensures that males always transmit unrecombined *Fat1^Flox^* alleles. This cross produces *Prx1-cre; Fat1^Flox/Flox^* mutants, and we used *Prx1-cre; Fat1^Flox/+^* mice as controls (occasionally *Fat1^Flox/Flox^*). To produce mice for the *Pdgfra-icre* cohort, there was no specific requirement the cre line to be provided in males or females, as there is no recombination in absence of Tamoxifen activation. *Pdgfra-icre; Fat1^Flox/+^; R26^YFP/+^* mice were crossed with *Fat1^Flox/Flox^; R26^YFP/YFP^* mice. Controls (referred to as *Ctrol^iPdgfra-YFP^*) include *Pdgfra-icre; Fat1^Flox/+^; R26^YFP/+^* and *Pdgfra-icre; Fat1^+/+^; R26^YFP/+^* mice. Genotyping was done as previously described for each line (Caruso *et al*., 2013; Helmbacher, 2018, 2022).

### Muscle lesions

For muscle injuries, mice were anesthetized by intraperitoneal injection of Ketamine (100mg/Kg) and Xylasine (10mg/Kg) and receive a subcutaneous injection of buprenorphin (0,1mg/kg). They were then placed on a heated pad, shaved around the area of interest, and disinfected with betadine. A small scalpel incision was made to open the skin above the target muscle, in order to displace the subcutaneous adipose tissue and visualize the muscle. This allows entering the needle in a well-defined position in the muscle, thereby also avoiding to damage large vessels. Lesions were made by intramuscular injections of glycerol (50% glycerol in sterile water), or Cardiotoxin (10 µM, dissolved in 0.9% NaCl solution). Injections were performed in the triceps brachii (TB) muscle in most cases, or occasionally in the tibialis anterior (TA) muscle, using an insulin syringe 0.3ml (30G). We injected a volume of 30 to 50 µl per muscle. Injections were always carried out in the left Triceps brachii, and the right side was used as uninjured control. After injection, the skin is sutured using Michel suture clips, applied with adequate clip-applying forceps (Fine science tools). Mice were kept under observation until they were fully awake and had recovered from the anesthesia.

### Tamoxifen induction

For the inducible model, CRE activation by Tamoxifen is carried out at adult stage. Tamoxifen (T5648, Sigma) is dissolved first in ethanol (100mg/ml) under agitation for 2h at 42°C, then diluted in Corn Oil (C8267, Sigma), at 10 mg/ml, kept under a chemical hood at room temperature, for the ethanol to evaporate, then aliquoted and stored at -20°C. Mice are injected intraperitoneally with a volume corresponding to 10µl/g (per gram of tissue weight) for mice weighing less than 30g (dosage 100µg/g). For mice weighing more than 30g, we limited the injected volume to 300 µl, to avoid liver toxicity of the Tamoxifen/oil mix. Each mouse (in both the control and mutant group) received 5 consecutive injections, with two injections before the lesion (day -2 and -1), and 3 injections after muscle lesion (days 0, +1, +2). The weight at first injection is considered as reference, and mice are weighed every day to evaluate potential weight loss. Any mouse having lost more than 15% after the two first injections is excluded from the experiment. Efficiency of recombination was assessed using the reporter line R26YFP, by immunohistochemistry with anti-YFP antibodies on muscle cryo-sections. Both *Ctrol^iPdgfra-YFP^* and *Fat1^iPdgfra-YFP^* mice carry one copy of the CRE allele and one copy of the R26-YFP allele, and have been exposed to the same dose and number of Tamoxifen injections, thus making *Fat1* the only varying factor.

### Tissue harvesting and immunohistochemistry

Muscles were collected after trans-cardiac perfusion of anesthetized mice with 4% PFA (in PBS). Mice were anesthetized with a lethal dose (2 to 5 fold the amount used for normal anesthesia) of Ketamine (200mg/Kg) and Xylasine (20mg/Kg) and received a subcutaneous injection of buprenorphin (0,1mg/kg). After perfusion, muscles were dissected and postfixed in 4% PFA (in PBS) for 4 to 6h at 4°C, rinced 3 times in PBS, and cryoprotected by immersion in 25% sucrose (in PBS), at 4°C until the tissues sank to the bottom (attesting equilibrium). Each muscle was transversally cut in two halves, the cut being made at the level of the center of the lesion. Tissue samples were embedded in gelatin/sucrose mix (7,5% gelatin, 15% sucrose, in PBS) in plastic molds. For embedding, the mix was preheated at 42°C, 2ml of gelatin/sucrose mix was poured in the molds. Excess liquid surrounding tissue fragments was dried on absorbing tissue, the fragments were deposited and oriented in the gelatin/sucrose mix, and the blocks were allowed to solidify on ice. A color mark indicating the axis and limits of tissue fragments was added on the solidified blocks prior to freezing in an isopentane bath cooled in dry ice ethanol mix. Samples were serially sectioned with a cryostat (10 µm sections), deposited on superfrost plus Gold microscope slides (K5800AMNZ72), which were stored at -20C. For immunocytochemistry, the slides were thawed, rinsed in PBS for 5 minutes, incubated with PBS 0,3% triton (P0.3T) for 5 minutes, then with 6% H202 in PBS 0,3% triton for 30 minutes, rinsed 3 times with P0.3T, and incubated overnight at 4°C with primary antibodies in blocking solution with 20% newborn calf serum, 0,3% triton in PBS. Slides are then washed by multiple incubations in with P0.3T under gentle agitation (minimum 5 x 20 minutes), and incubated in blocking solution (as above) with secondary antibodies conjugated with fluorophores, either overnight at 4° or at room temperature for at least 1h30. After multiple washes with P0.3T, the slides are mounted under coverslips with mounting medium with ProlongGold antifade reagent and DAPI (P36935, Thermo Fisher Scientific), and kept at 4°C. The list of primary antibodies used is provided below. Images are acquired using a Zeiss Z1 microscope, equipped with an Apotome, using the Zen software. After initial treatment with Zen, final images are exported in tif, and treated with ImageJ and FiJi for image analyses and with Photoshop for figure mounting.

Primary antibodies and reagents: Chicken anti-β-galactosidase, Abcam ab9361, RRID: AB_307210. Purified rat anti-mouse CD104a (Pdgfra) monoclonal antibody, clone APA5, BD Biosciences #558774, RRID: AB_397117; XP Rabbit anti-Perilipin monoclonal antibody, clone D1D8; Cell signaling technology, #9349, RRID: AB_10829911; Chicken anti-GFP, Aves, #GFP-1020, RRID: AB_10000240; Rat monoclonal anti mouse Tenascin-C (TnC) antibody, clone Mtn-12, Thermofisher Scientific, #MA1-26778, RRID: AB_2256026; Mouse monoclonal anti-Chicken PAX7, IgG1, Developmental studies hybridoma Bank, #Pax7, RRID: AB_528428; Rabbit anti-Iba1 polyclonal antibody, Wako Chemicals, #019-19741, RRID: AB_839504; Alexa-Fluor-conjugated Phalloidin, Thermofisher Scientific, Alexa-Fluor-488: A12379; Alexa-Fluor-546: A22283; Alexa-Fluor-647: A22287.

### β-galactosidase assays

Staining for β-galactosidase activity on cryosections was done as described previously (Caruso *et al*, 2014; Helmbacher, 2018), using Salmon Gal (6-chloro-3-indolyl-β-D-galactopyranoside from Appolo scientific, Manchester, UK) as substrate. Briefly, slides with cryostat sections were thawed in PBS, incubated overnight at 37°C in a solution containing 4 mM Potassium Ferricyanide, 4 mM Postassium Ferrocyanide, 4 mM MgCl2, and 0,04% NP40 substitute, without substrate, to inactivate endogenous enzymes, rinsed 3 times in PBS, and incubated in a solution with Salmon Gal (1mg/ml), nitro blue tetrazolium (NBT, N6876 Sigma, 330 µg/ml), 2 mM MgCl2, 0,04% NP40 substitute, in PBS. Staining intensity was monitored under a dissection microscope. If the solution became purple, it was replaced with fresh staining solution until completion. After staining (1-2h for sections of *Fat1^LacZ^* muscles) the slides were fixed overnight with 4% PFA in PBS, and mounted.

### Image analyses and quantifications

For every mouse included in the experimental cohorts, muscle samples were serially cryo-sectioned, so as to generate a collection of representative slides to use for multiple immunohistochemistry experiments. This allowed performing multiple assays on the full cohort. After performing a given IHC experiment, we acquired a 10X mosaic image of at least one slice containing the entire muscle section. Quantifications were systematically done on 10X images of the entire muscle section rather than portions of it acquired at higher resolution to avoid selecting areas of interest in a biased manner. Instead, we proceeded in a completely unbiased manner, following a systematic set of steps described below:

All IHC included a channel with phalloidin staining. Single channel TIF images exported from ZEN were assembled as distinct layers in Photoshop, with fusion modes for each layer set to “lighten”, on top of a background layer filled with black. Each layer was duplicated so as to maintain unmodified images, while modifications were done on duplicates. An extra duplication of the phalloidin layer was done with enhanced midtones, so as to visualize all fibers equally well irrespective of phalloidin intensity. The lasso tool was used to manually select areas containing undamaged fibers, and the content of selected areas was deleted in all (duplicate) layers. After completing the step by step deletion of all undamaged fibers, each remaining content of each layer was individually merged with the black background layer, thus generating for each color a tif document to be analyzed in FiJi/ImageJ, containing the staining corresponding, for each color/antibody, to the content of the lesion. These documents were opened in ImageJ, converted to 8bit, and thresholded (using the same thresholding parameters and exposure time for all compared images). The area of the objects contained in each image were analyzed using the “analyze particles” tool (not including holes) and exported to excel. The area, measured in square pixels, was converted in real dimension by multiplying the object area by multiplying with the square resolution (96 ppi, pixels per inch), and with the square pixel size corresponding to the magnification used (0,65 for 10x images, and 0,32 for 20X images). Area (um2) = Area (px2)x(96)^2^x(pixel size)^2^. For each image/sample/antibody-color, the objects measured were ranked by size (from larger to smaller), a fixed area cutoff (of 50µm^2^) was applied to delete artefacts smaller than the cutoff size. For analyses of PLIN1, TnC, YFP, stainings, the total area of remaining objects was then calculated, and expressed as raw area or relative to the lesion area (measured in axiovision software). For analyses of fiber cross section areas, we defined ranges of 300 µm^2^, and counted the percentage of fibers in each size range. For analyses of PLIN/YFP double staining, the YFP-positive layer was divided in PLIN+ or PLIN1-negative, using the color selection tool, to selectively delete PLIN+ cells or Plin1-negative cells, from the YFP+ layer, thus generating two distinct images, in which we applied the method described above (threshold, particle analyses, cutoff, sum), to quantify for each sample the total PLIN1+;YFP+ area, and the total PLIN1-negative;YFP+ areas. For analyses of PDGFRA/YFP double staining, this was done manually, by scanning the image for YFP+ cells, and for each cell, defining it as PDGFRA-negative, intermediate, or high.

Quantification of signal intensities in Figure S1D were performed using Fiji as previously described (Caruso *et al*., 2014; Fan *et al*, 2015). Briefly, the area shown in Figure S1C was cropped from larger images, to span an area going from the uninjured muscle to the center of the lesion, where degenerating fibers undergo necrosis. Each color channel was opened in Fiji and converted to 8bit images. Staining intensity was measured using the function > analyze > Plot profile, along an area matching the whole length of the cropped image, and a few pixel in height.

This measurement was done 4 times for each color (on windows of full width, and a few pixels heights). The plot values were subjected to background subtraction, threshold subtraction (to avoid negative values), and expressed as a percentage of the max amplitude (defined as the difference between max intensity and background intensity; the max intensity value being defined as the average value in a manually defined window matching the active zone.

### Statistical analyses

All measurements above yielded one single numerical value per mouse, allowing to have in each cohort, and for each genotype, sufficient numbers of mice to perform comparisons and statistical analyses. Sample size corresponds to the number of independent animal in each group/genotype/stage. No specific method was used to define sample size. As we did not know prior to initiating, whether the phenotypes were the same irrespective of sex or if there were sex-specific phenotypes, we included in each cohort sufficient (and equal) numbers of males and females, so as to perform statistics on separated sexes and conclude reliably on sex as parameter. Results are presented either by pooling males and females, or by separating them (supplementary figures 5 and 8). All statistics tests were done on comparisons between two genotypes, using unpaired Student’s t-Test, when the data showed a normal distribution and equal variance between the two groups, and Mann-Whitney test otherwise. Differences were considered significant when p<0,05. All p-values are indicated in the figures, unless the comparison yielded a radically non-significant result (in which case we indicated ns).

## Data availability

The datasets supporting the conclusions of this article are included within the article and its additional files, with the exception of figures 1C, and 1H, and Figure 2, for which we used the following published gene expression datasets. Data on Fat1 expression after CTX and Glycerol injury (Figure 1C) were extracted from ref (Lukjanenko *et al*., 2013), and are available in the Gene Expression Omnibus Repository, under the accession number GSE45577. Data on gene expression in biopsies from human muscular dystrophy patients (Figure 1G) were extracted from references (Bakay *et al*., 2006; Dadgar *et al*., 2014), and are available in the Gene Expression Omnibus Repository under the accession number GDS1956, or GSE3307. ScRNAseq data used in Figure 2 were extracted from the ScMuscle dataset, a set of ScRNAseq data integrated in a common visualization tool published in ref (McKellar *et al*., 2021), initially available as a web application (scmuscle.bme.cornell.edu) or currently available as fully processed Seurat and CellChat objects that can be downloaded from the datadryad platform (doi: 10.5061/dryad.t4b8gtj34). The full list of accession numbers, including a novel dataset GSE162172 (transcriptomic sampling of regenerating aged mouse hindlimb muscle after notexin injury), as well as several previously published datasets, is available in the supplementary Table 1 of ref (McKellar *et al*., 2021).

## Acknowledgements

We thank Flavio Maina and Robert Kelly for helpful discussions and suggestions, Osvaldo Contreras for critical reading and improvement of the manuscript, Dominique Fragano for mouse genotyping, the IBDM mouse facility staff for animal housing and care, Charline Ytier and Clara David for their help in setting up the new lab, and for critical reading of the manuscript. We thank the IBDM direction for temporary supporting animal housing charges, and Frederique Magdinier for help in our recent lab move and for providing a welcoming and supportive scientific environment.

## Author information

### Authors and Affiliations

Aix Marseille Univ, CNRS, IBDM UMR 7288, Parc Scientifique de Luminy, Case 907, 13288 Marseille, France;

Pierre-Antoine Ferracci, Françoise Helmbacher.

Aix Marseille Univ, INSERM, Marseille Medical Genetics, U1251, 13005 Marseille, France Françoise Helmbacher

## Contributions

**Pierre-Antoine Ferracci :** investigation, data analysis, manuscript reviewing and editing. **Françoise Helmbacher :** Conception, investigation, data analysis, mouse experimentation, funding acquisition, supervision, figure assembly, manuscript original draft writing, reviewing and editing.

---

**Figure S1:**
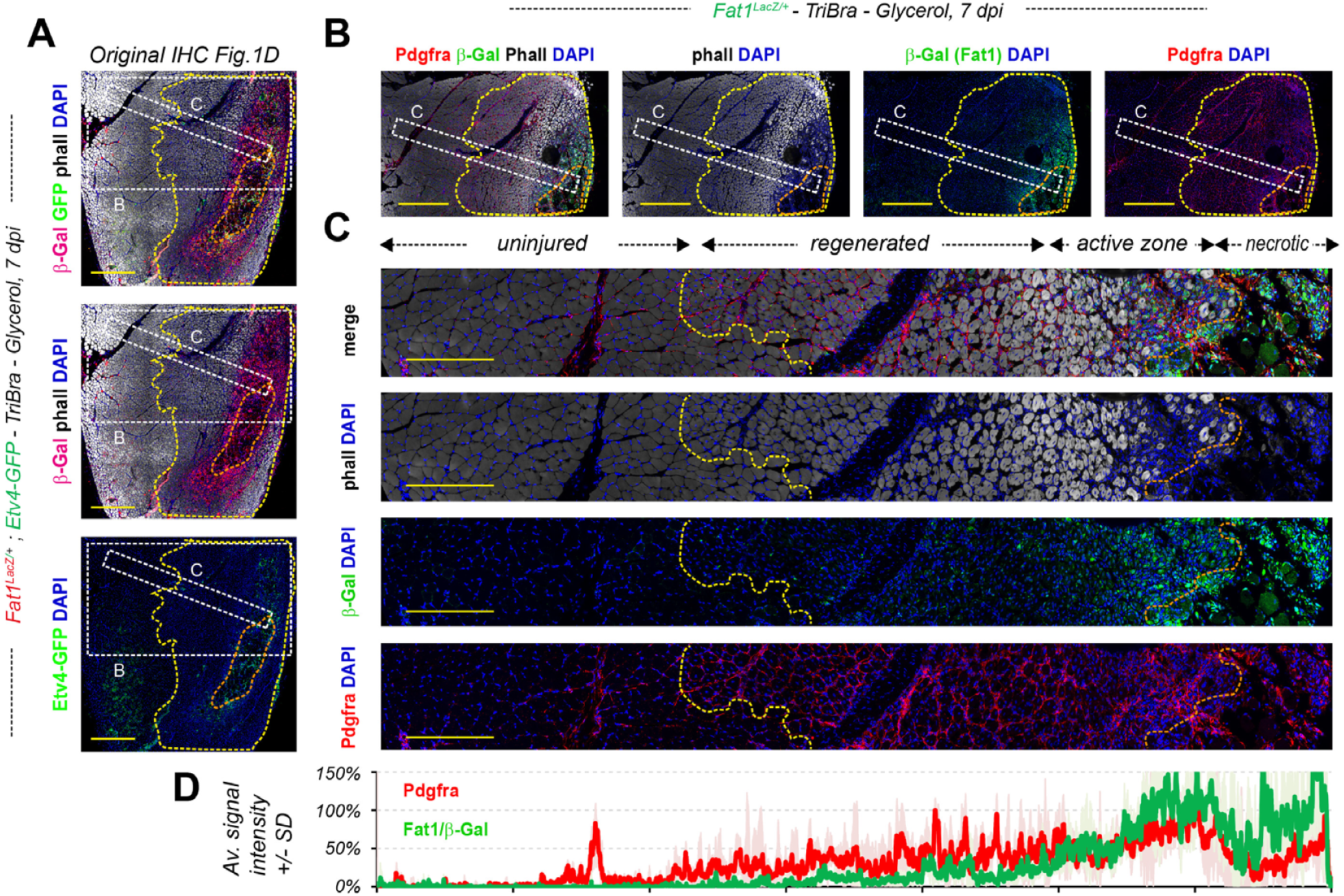
Fat1 expression is transiently induced in FAPs and myogenic cells in regenerating muscle after injury. (A) The IHC experiment shown in Figure 1D is presented here in its original setting. The sample was a Triceps Brachii muscle from a *Fat1^LacZ/+^; Etv4-GFP^+^* mouse, injured with glycerol, and collected at 7dpi. The section was immunostained with antibodies against beta-galactosidase (here in Red), and against GFP (Green, highlighting Etv4-GFP expression), combined with Alexa-conjugated phalloidin (white). The images presented in Figure 1D were color converted for simplicity of the message. (B, C) From the same animal, additional IHC were performed on alternate sections of the same muscle, with antibodies against beta-galactosidase (here in Green), and against Pdgfra (Red), combined with Alexa-conjugated phalloidin (white). Acquisitions at the scale equivalent to those in (A) were not done, but we acquired images with 20X magnification, shown full size in (B), and at higher magnification in Figure 1E (crops of defined size). An alternative way of showing how expression levels vary across different areas of the section is shown here in (C), representing crops of a band the size of which is indicated as white dotted lines in (A). (D) quantification of signal intensities for Fat1/β-Gal (green) and Pdgfra (red) along images in (C): each plot represents the average intensity (+/- standard deviation in light green and light red, respectively) as percentage of Max, for 4 narrow bands along the images. Scalebars: (A) 500 µm; (B) 500 µm; (C) 200 µm.

**Figure S2:**
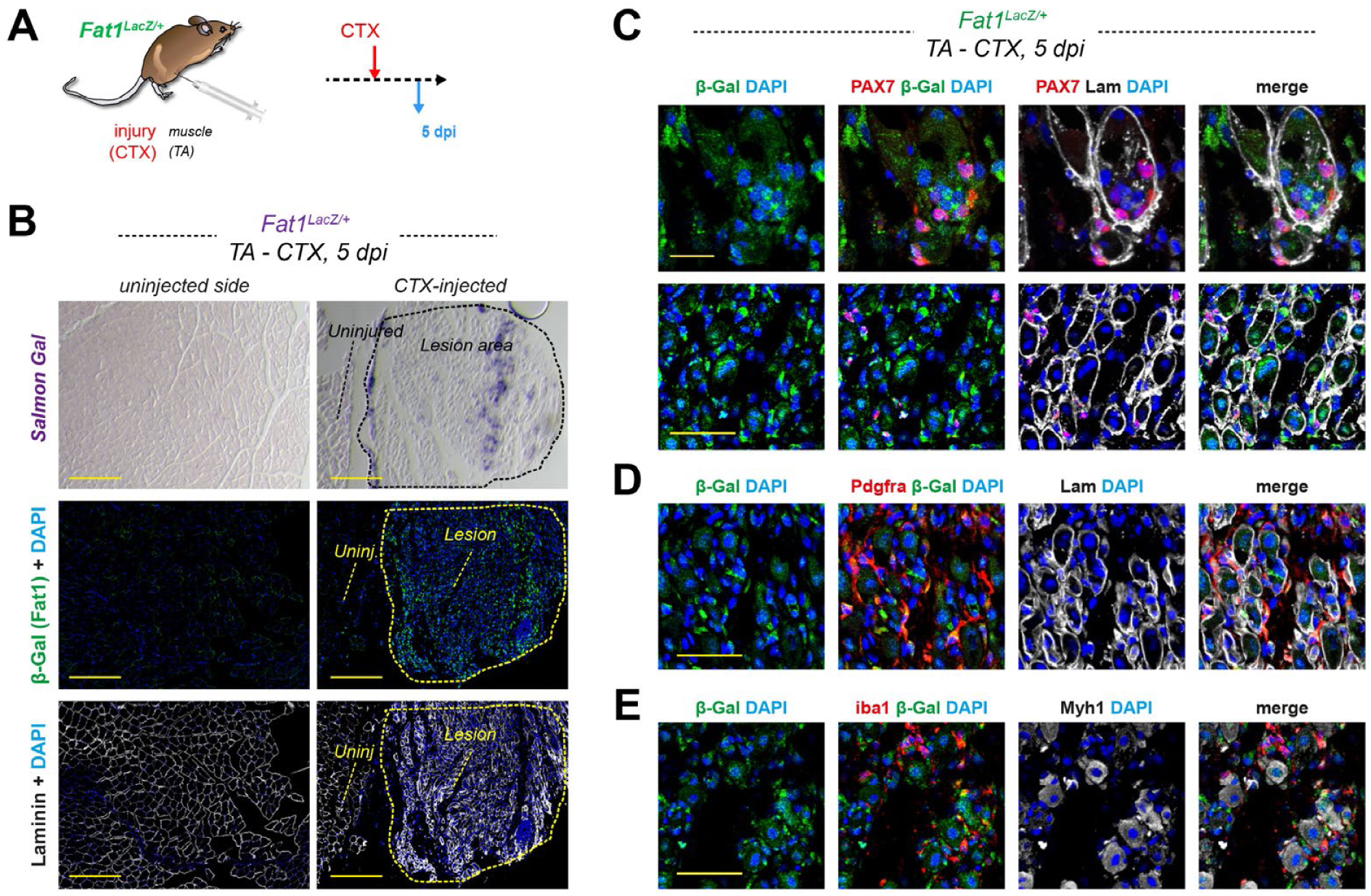
***Fat1^LacZ^*induction by Cardiotoxin (CTX), 5 days after injury .** (A) Scheme of the experiment, in which adult *Fat1^LacZ/+^* mice were injured by injection of CTX in the Tibialis anterior muscle (TA), and showing the timeline, with injury occurring at day 0, and muscle collection for histological analyses at 5 days post-injury. (B) *Fat1^LacZ^* expression is visualized on the same muscle cross-sections as in Figure 1B of a *Fat1^LacZ/+^* mouse 5 days after CTX injury in the TA, with unijected (left) and injected (right) sides, by Salmon gal staining (top), or immunohistochemistry with anti-β-galactosidase (green) and laminin (white) antibodies (with DAPI, blue), showing that expression is undetectable in the uninjured adult muscle, but induced by the CTX injury. (C, D, E) high magnification images at the level of the actively regenerating area of the lesion, to show at different magnifications the combination of anti-β-galactosidase (green) and laminin (white) with Pax7 (red, C), Pdgfra (red, D, and iba1 (red, E). Scalebars: (B) 500 µm; (C) top images: 20 µm, bottom images 50 µm, (D,E) 50 µm.

**Figure S3:**
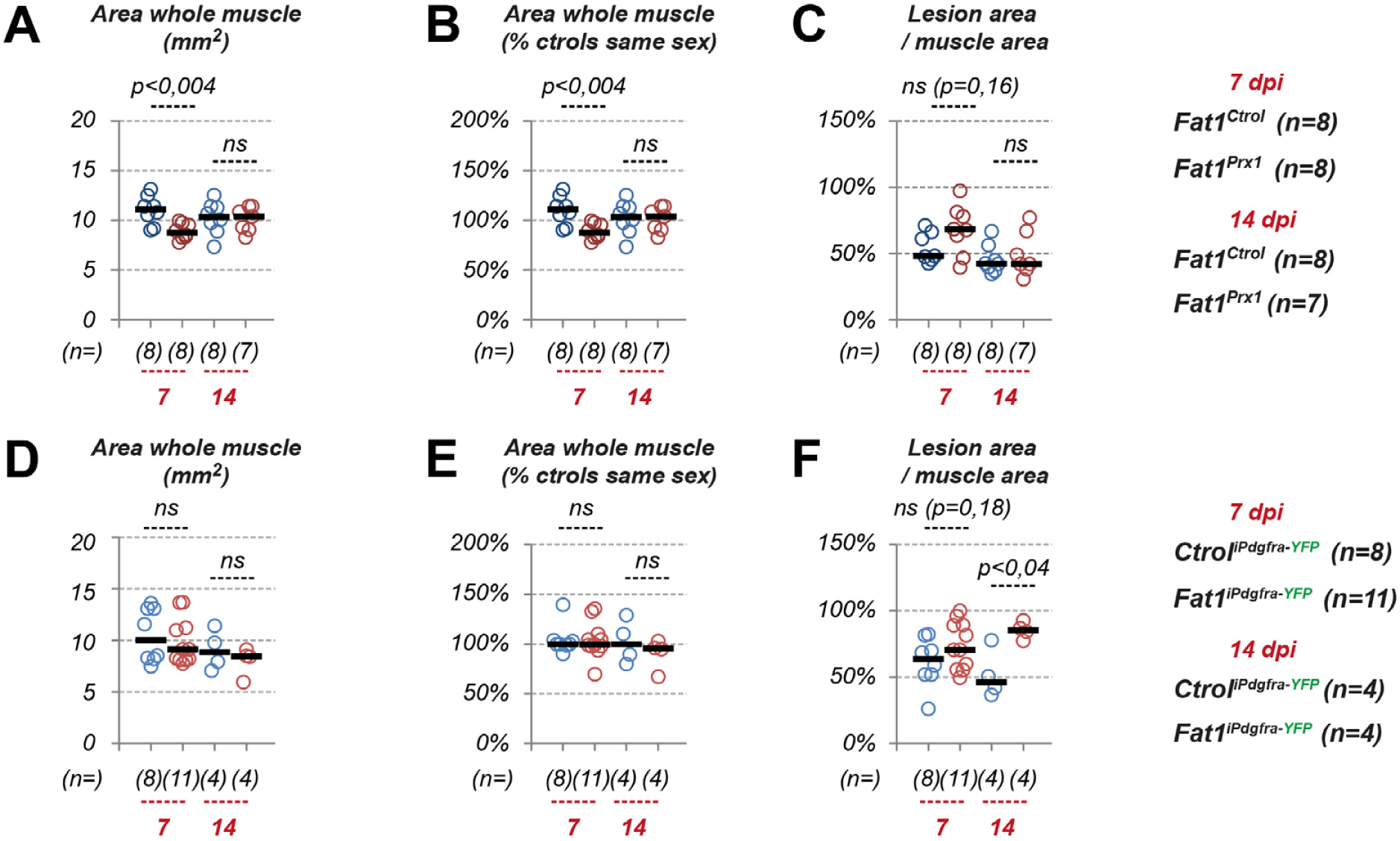
Presence and absence of pre-injury phenotypes in the Prx1-cre and inducible Pdgfra-icre models. Quantifications of (A, D), the whole muscle area (in mm2), (B, E) the whole muscle area (normalized to the median area of controls of the same sex), and of (C, F) the ratio between lesion area and whole muscle area, in *Fat1^Ctrol^*and *Fat1^Prx1^* mice (A, B, C), and in *Ctrol^iPdgfra-YFP^*, and *Fat1^iPdgfra-YFP^* mice treated with Tamoxifen as described in Figure 5B (D, E, F) showing individual data points for each mouse, at 7 and 14 dpi. The reduction in muscle area observed in *Fat1^Prx1^* mice at 7dpi, associated with an increased L/M ratio (not significant), indicates that a phenotype specific to the uninjured part of the muscle pre-existed the lesion. This phenotype is transient, and no longer detected at 14 dpi, indicating that myogenic repair in the lesion rescues the muscle mass equally well in *Fat1^Ctrol^* and *Fat1^Prx1^*mice. In contrast, in the inducible *Pdgfra-icre* model, the lack of reduction in whole muscle area shows that there are no pre-injury phenotype before Tamoxifen treatment. This allows uncovering an effect on lesion size at 14dpi, potentially accounted for by enhanced adipogenesis in *Fat1^iPdgfra-YFP^* mice.

